# Toward a Random Background for Ligand Optimization

**DOI:** 10.64898/2026.05.10.724162

**Authors:** Xinyu Xu, Olivier Mailhot, Galen J. Correy, XP Huang, Joao Braz, Da Shi, Karthik Srinivasan, Kara Zielinski, Yuliia Holota, Yuliia Kuziv, Christos Tsoutsouvas, Nathan Levinzon, Yagmur U. Doruk, Moira Rachman, Morgan Diolaiti, Maisie Stevens, Fangyu Liu, Katie Holland, Harald Hübner, Jing Wang, Yujin Wu, Alan Ashworth, Alexander Makriyannis, Yuqi Zhang, Yurii Moroz, Peter Gmeiner, Robert Abel, Aashish Manglik, Allan I. Basbaum, Bryan L. Roth, James S. Fraser, Brian K. Shoichet

## Abstract

Ligand optimization is central to drug discovery as hundreds of analogs might be designed and synthesized between an initial hit and a therapeutic candidate. The efficiency of this process is unclear, at least partly because there is no random background for optimization against which to compare. Such a random background might emerge from synthetically accessible but otherwise systematic random small substitutions across starting ligands, measuring likelihood of achieving a substantial improvement in affinity/potency or other property by any single perturbation. Recent literature and ligand-affinity/potency databases suggest that perhaps 10% of analogs with minor modifications improve upon a parent’s potency substantially (by ≥10-fold), but this number is clouded by reporting bias, intentional improvement, and inter-group reproducibility. To begin to establish a background expectation for ligand optimization, we comprehensively and systematically modified 18 lead molecules across six targets with single atom changes; 257 compounds were synthesized. Unexpectedly, 11.2% of these random small perturbation analogs improved potency by ≥10-fold over their parents. Conversely, these more potent analogs typically had worse *in vitro* pharmacokinetics (e.g. reduced metabolic stability, lower plasma free fraction). While it was possible to find analogs where the potency increase compensated for inferior exposure and half-life, resulting in more potent compounds in vivo, overall a frustrated landscape for ligand optimization is revealed. This study begins to establish a background expectation for ligand potency optimization and offers a simple strategy to do so. It also begins to quantify the challenges confronting the field in moving beyond in vitro potency.

## Introduction

Ligand optimization is central to chemical probe and drug discovery, as initial active molecules rarely have the potency or pharmacokinetic properties to be viable in vivo^1,2^. Accordingly, between the discovery of an initial hit and a clinical candidate many hundreds, occasionally thousands of optimized analogs might be synthesized^1,3^. Several strategies^4–6^ ^7,8^ have been developed to improve this process, ranging from early empirical approaches such as Topliss Trees^9^ and Hansch QSAR^10^ to contemporary AI-guided ADMET^11^ prediction and free energy calculations^12,13^. How efficient these strategies are remains uncertain, as there is no random background against which to compare them. If we use a ten-fold improvement between analog and parent as a benchmark for substantial impact, then only about 10% of analogs meet this standard in the ChEMBL ligand-protein activity database (**Extended Data Figure 1** a)^14^. Since these ChEMBL results suffer from success bias, sample multiple types of perturbation, and struggle with inter-group irreproducibility^15^, the public domain offers no sure guidance on what level of improvement one might expect in ligand optimization if one were making random conservative changes.

Random backgrounds have long been used in biology to help quantify significance. In genomics, they help to distinguish between artificial and natural selection^16^. In epidemiology, random incidence rates help distinguish genetic diseases from those driven by environmental factors^17,18^. In protein engineering, random backgrounds are used to evaluate improvements in enzyme function^19^, and alanine scanning has been used to introduce a minimally-biased perturbation to find hot spots for ligand binding and protein function^20^. A random background for ligand optimization, conceivably, could quantify progress across chemotypes and targets and compare different optimization strategies.

Akin to alanine scanning, an unbiased background for ligand optimization might involve conservative perturbations, systematically and comprehensively applied across multiple unrelated parents across several different targets. Muegge and colleagues have described a “positional analog scanning” approach, which substitutes ligand non-polar hydrogens (C—Hs) with groups like CH₃, OH, Cl, F, and Br, and aromatic carbon atoms with nitrogen^21^. In retrospective analyses of the ChEMBL24 database^22,23^, effects on affinity, functional activity, solubility, clearance, and permeability were studied. Among over 110,000 matched molecular pairs (MMPs), about 30% of the time of one analog improved over 3-fold in affinity/potency versus the other. Our understanding of this observation is tempered by design and reporting biases toward compound improvement, by the variability in experimental conditions across datasets that can reduce reproducibility^15^, and by the few parent compounds that had multiple conservative changes. Thus, it seemed interesting to create a set of parent ligands that were systematically and comprehensively modified by small perturbations, without design and without a bias toward improvement, using the positional analog scanning approach^21,24^ where the differences between parent and the suite of analogs were quantified by a single lab.

To create what we will refer to as a random background set for ligand optimization, we chose 18 parent ligands spanning six targets, including three GPCRs (alpha2A adrenergic receptor (α2a), μ-opioid receptor (MOR) and cannabinoid receptor type 2 (CB2)), one transporter (the serotonin transporter, SERT) and two soluble enzymes (AmpC β-lactamase (AmpC), and Macrodomain 1 of SARS-CoV-2 (Mac1)). This set was chosen for targets that we had under experimental control and is admittedly far from comprehensive: we explore only class A GPCRs, no ion channels nor kinases are represented, only one transporter, and only two of almost innumerable soluble enzymes. The 18 parent compounds were chosen for similar pragmatic reasons—they were molecules to which we had ready access. Still, these proteins provide a range of target classes, while the 18 parents cover a wide range of physico-chemical properties (**Extended Data Table 2**) and values, ranging from 1 nM to 43 µM (**Table 1**), and most were topologically unrelated. For each of the 18 parents we systematically explored all analogs accessible by a single atom modification where such modifications were synthetically feasible—typically costing no more than $400 to acquire—and where they did not modify the net charge of the parent at physiological pH. The single atom modifications involved substituting carbon hydrogens with one of methyl^25^, hydroxyl^26^, chloro^27^, fluoro^28^, or rarely bromo, or substituting aromatic carbon atoms with nitrogens (**Figure 1**). These changes are not comprehensive, but they do represent changes preferred by medicinal chemists while avoiding those that would introduce oxidation liabilities (e.g., adding a thiol or aromatic sulfur) or changes of valence (e.g., nitrogen to oxygen or sulfur); moreover, they were made without design and as systematically as pragmatism would allow.

**Fig. 1:**
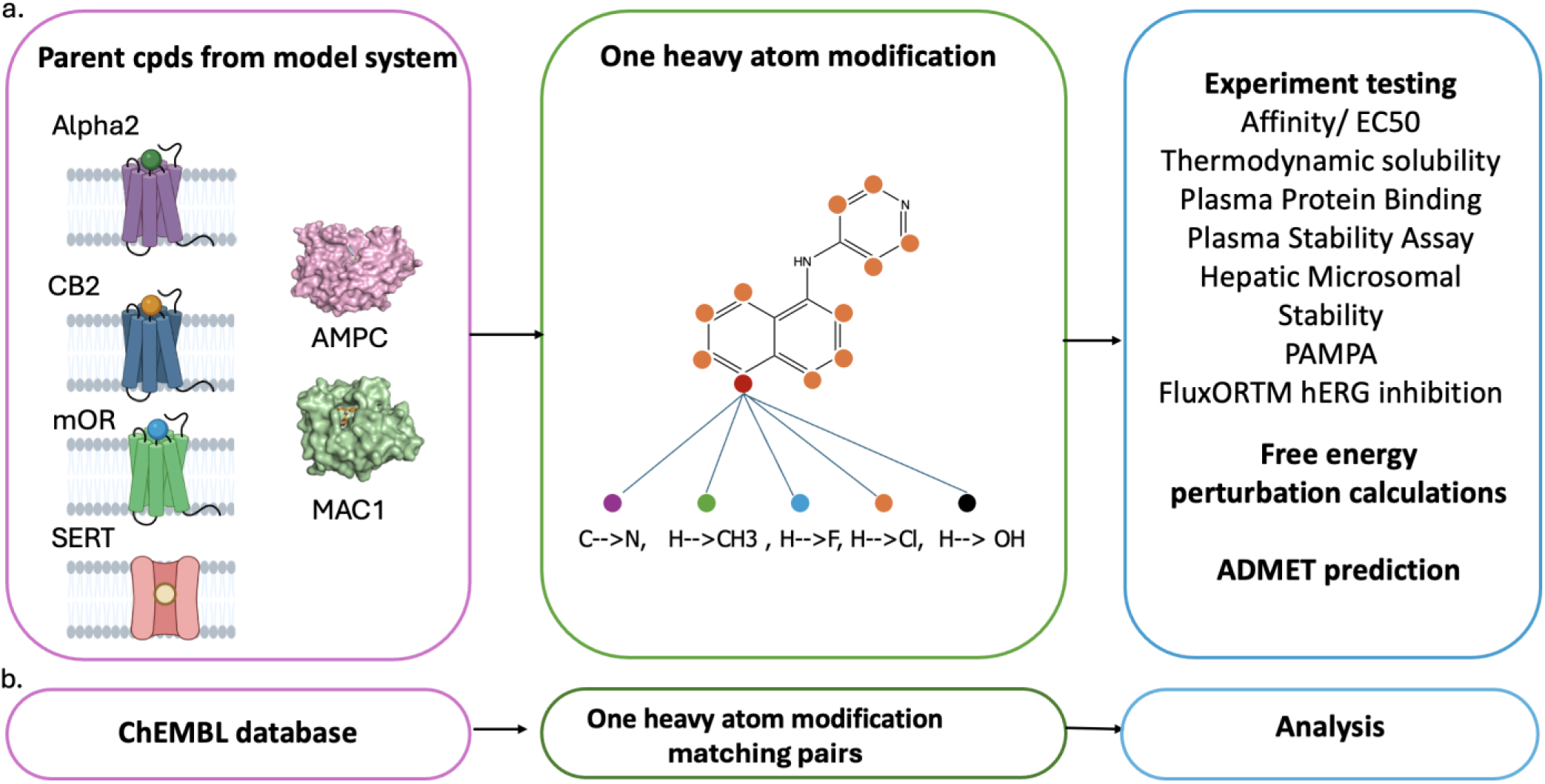
Experiment and Analysis Workflow for One Heavy Atom Analogs. **a**, Parent ligands from six drug targets (the aminergic GPCR Alpha2a-AR, the lipid GPCR CB2R, the peptide GPCR µOR, the serotonin SLC transporter SERT, and the soluble enzymes AmpC and Mac1) were modified by one-at-a-time, one atom substitutions: H→CH3, H→Cl, H→OH, H→F, and ring C→N. These modifications were made systematically and comprehensively; only compounds that were too expensive to synthesize, or that changed the charge state of the parent were left out. The analogs were tested for changes in affinity/potency, solubility, plasma protein binding, plasma stability, hepatic microsomal stability, permeability, and hERG inhibition, versus their parent ligands.

**Table 1.** Analogs from all parents improving affinity/potency by ≥3x or ≥10x.

| Protein | Assay readout | Parent Ligand | Parent Affinity/potency | Total Analogs | Analogs with $\geq 3$X Affinity/potency improvement | Analogs with $\geq 10$X Affinity/potency improvement |
| --- | --- | --- | --- | --- | --- | --- |
| MOR | EC50 | Z4407498716 | 9.2 nM | 16 | 2 | 2 |
| alpha2 | EC50 | Z2750653629 | 1.4 nM | 31 | 7 | 4<sup>a</sup> |
| alpha2 | Ki | Z3034773248 | 22.1 μM | 18 | 3 | 1 |
| alpha2 | EC50 | Z4376630014 | 8.3 nM | 10 | 2 | 0 |
| alpha2 | Ki | Z8727395870 | 13 μM | 18 | 1 | 0 |
| AmpC | Ki | Z2610488449 | 24.2 μM | 10 | 2 | 0 |
| cb2 | Ki | Z52076138 | 2.3 μM | 7 | 7 | 6<sup>b</sup> |
| cb2 | Ki | Z6969215903 | 0.4 μM | 6 | 0 | 0 |
| cb2 | EC50 | Z8184698918 | 0.1 μM | 20 | 2 | 2 |
| Mac1 | IC50 | Z5398393122 | 2.2 μM | 5 | 0 | 0 |
| Mac1 | IC50 | Z1039063794 | 0.7 μM | 12 | 0 | 0 |
| Mac1 | IC50 | Z7534253453 | 8.9 μM | 22 | 8 | 2 |
| SERT | Ki | Z2009978218 | 1.1 μM | 14 | 4 | 0 |
| SERT | Ki | Z2573292480 | 0.08 μM | 11 | 1 | 0 |
| SERT | Ki | Z2573292509 | 0.2 μM | 9 | 2 | 1 |
| SERT | Ki | Z6971277399 | 13.4 μM | 15 | 10 | 4 |
| SERT | Ki | Z8727393896 | 42.7 μM | 14 | 8 | 3<sup>c</sup> |
| SERT | Ki | Z8731642686 | 1.8 μM | 19 | 10 | 4 |
| Total |  | All Parents |  | 257 | 69 | 29 |

\*Fold-changes are displayed as nearest-integer values; exact (unrounded) fold-changes are provided in Supplementary Table S1. Using exact fold-changes, 62 analogs improve $\geq 3$ -fold and 25 improve $\geq 10$ -fold. **a.** One of these had a fold change of 9.7. **b.** Two of these had fold-changes of 9.7 and 9.8. **c.** One of these had a fold-change of 9.9, see Supplementary Table S1.

These substitutions were systematically and comprehensively applied without design, ensuring an approach unbiased by anything other than synthetic accessibility and price (which we admit are meaningful constraints, as we make clear below). Overall, 257 single-atom analogs were synthesized and tested for changes in activity on the target, allowing us to measure how often we might expect affinity/potency to improve ≥10-fold by minimal perturbation. This begins to provide a background expectation for how often a *designed* analog might meaningfully improve affinity/potency. As we will show, these undesigned changes had an unexpectedly high success rate, suggesting also a systematic strategy for optimization, even though that was not our primary goal. Since affinity/potency maturation is only one criterion for advancing from hit to candidate, we also measured in vitro pharmacokinetic properties for the analogs, including permeability, metabolic stability, plasma protein binding, and three other terms, versus the parent compounds. These pharmacokinetic studies explore what are perhaps inevitable trade-offs between pharmacokinetics and target affinity/potency in ligand optimization.

## RESULTS

We began with 18 parent ligands, most of which are unrelated to one another (ECFP4 Tanimoto coefficients, Tcs, <0.35), with affinities ranging from 1 nM to 43 µM and molecular weights (MW) ranging from 200 to 400 amu (**Extended Data Figure 2**). Each of the 18 parents was modified, one atom at a time, with five types of single atom substitutions (**Figure 1a**, central panel), subject only to the expense and charge modification constraints (above). About 7% of the possible molecules met these criteria and of those ∼80% were successfully synthesized, resulting in 257 analogs. In the analyses that follow, we pool all 257 analogs for analysis, which reduces the noise from a single scaffold or single target and improves statistical power (**Supplementary Figure 1**).

These 257 analogs were tested for activity changes (K_d_, IC_50_, or EC_50_) versus the parent ligands across the six targets. For the μ-opioid receptor, EC_50_ values were determined in cAMP Glo-sensor assays. α2A and CB2 ligands were assessed by cAMP assays and by radioligand competition binding with ^3^H-rauwolscine or [³H]CP-55940, respectively. Serotonin transporter interactions were analyzed using [^3^H]-citalopram binding assays. AmpC β-lactamase (AmpC) inhibition was measured enzymatically while that of Mac1 was measured by ligand displacement. For every parent, and for most receptors overall, we only report changes for one type of activity, i.e., if a parent K_i_ is reported than only K_i_ values are reported for its analogs, and if a parent is an agonist, only EC_50_s are reported for its analogs. Each functional or inhibition assay is done in reference to a literature positive control molecule (**Supplementary Figure 2**). Representative concentration–response curves and the corresponding Z values for each assay are provided in **(Supplementary Figure 3)**. Especially for agonists, the effect of receptor expression is controlled for **(Supplementary Figure 4, 5)**. Because changes in agonist efficacy can sometimes affect interpretation of EC_50_, we inspect efficacy values to ensure that they are similar between parent and analog; full concentration-response curves and analyses are shown for every parent-analog pair, allowing for direct inspection of these results.

Overall, 23% of the analogs affected activity by ≤2-fold (fold changes rounded to the nearest integer) (**Supplementary Table 1, Supplementary Figure 1**), while 30% *reduced* activity by ≥10-fold **(Extended Data Figure 1** b). Intriguingly, of the 257 analogs tested, 29, or 11.3%, improved activity by ≥10-fold rounding to the nearest integer (**Table 1**). We note that 4 of these 29 had fold-changes of between 9.7 and 9.9 to 1 decimal place; we consider these effectively 10-fold improvements and used the same convention in analyzing the public ChEMBL data (see **Table 1** and **Supplementary Table S1** for details). Such ≥10-fold improvement analogs were found for five of the six targets, and for 10 the 18 parents (**Table 1**). The effects and chemical identities of all analogs relative to their parents are listed in **Supplementary Table 1**.

We were interested in which perturbations had the biggest effects, how the rates of affinity/potency improvements compared to the literature, with its admitted biases, and if physical properties of the parent compounds or of the binding sites correlated with greater likelihoods of affinity/potency improvements. Among the analogs, methyl substitutions were the most likely to improve activity ≥10-fold, with chlorine substitutions the second most likely; for ≥3-fold improvements, of which there were 69 among the 257 analogs, both chlorine and methyl substitutions were the most common (**Figure 2a**). Given the similar size and similar hydrophobicity of methyls and chlorines, their similar effects may be rationalized. Conversely, fluorine substitutions, which are often used to increase metabolic stability, yielded no ≥10-fold and few ≥3-fold improvements in activity. Aromatic carbon to nitrogen substitutions also rarely improved affinity/potency, though as we will see they often had the most favorable impacts on pharmacokinetic properties.

**Fig. 2.**
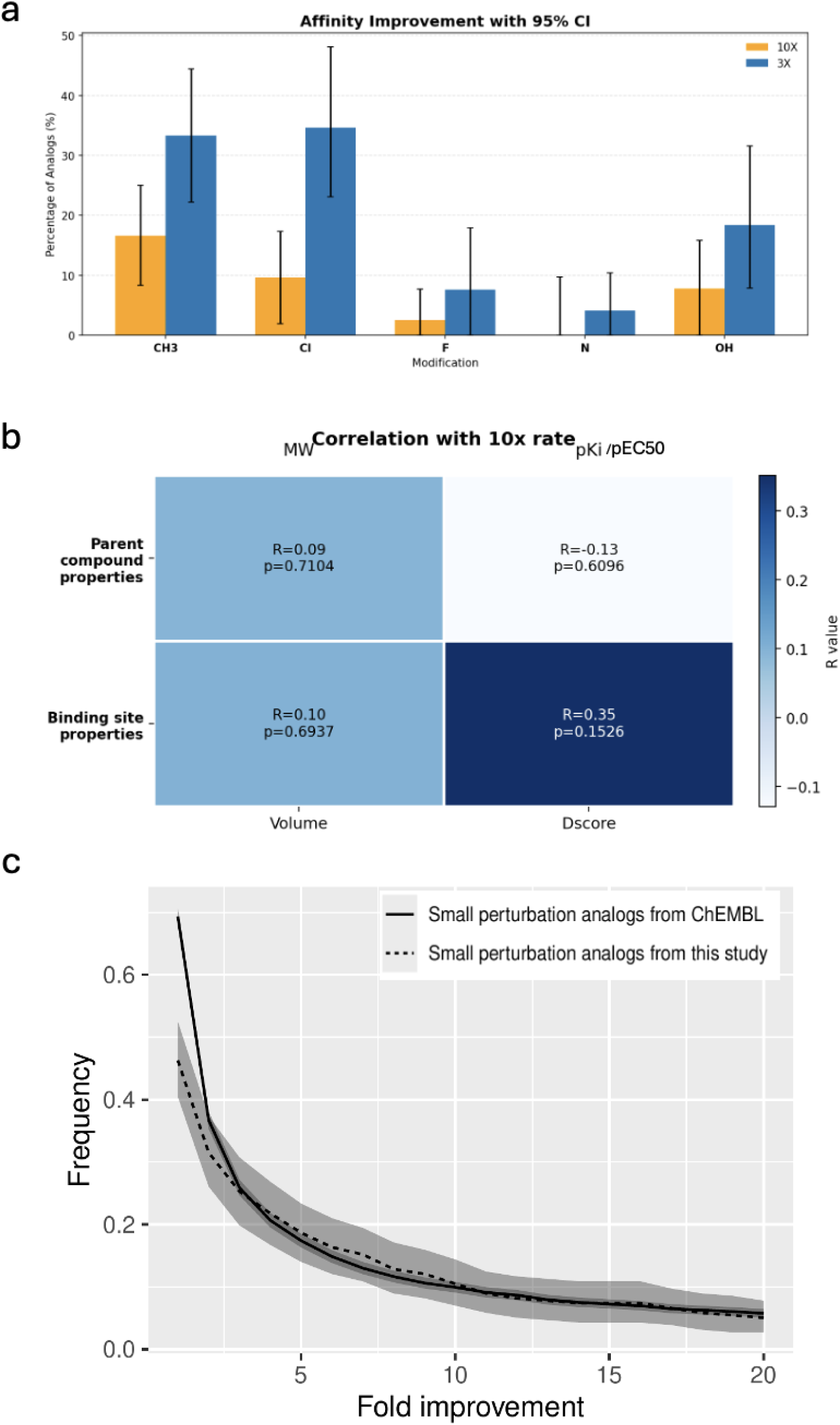
Property correlations of affinity/potency improvement and comparison with literature values. **a.** The fraction of compounds improving by ≥3- or ≥10-fold in activity by type of single modification (CH₃, F, Cl, Br, N, OH). **b**. Correlation heatmap between physicochemical properties of binding pockets and parent compounds with log₁₀ of affinity/potency fold-change. **c**. The fraction of single atom modifications (CH₃, F, Cl, N, OH) that improve activity, comparing the experimental observations in this study with results from the ChEMBL database. Shaded regions indicate 95% confidence intervals, estimated from 10,000 iterations of bootstrap resampling. The ChEMBL ligands are a composite of 54,021 parents and 79,984 small change analogs; for only 4.7% of these were there more than three analogs per parent.

Because increases in hydrophobicity will increase affinity/potency simply by disfavoring the unbound state, and have physical property liabilities, it is useful to consider the improved affinity/potency of the analogs in light of ligand efficiency (LE) and liphophilic efficiency (LiPe), which control for these effects. If affinity/potency improvements were largely driven by hydrophobicity we would expect to see both terms deteriorate for the analogs. Instead, for analogs that improved ≥10-fold both ligand efficiency and lipophilic efficiency much improved.

For instance, an analog with an added methyl, chloro or other atom that improved in binding or in EC_50_ by even 3-fold enjoyed a ligand efficiency of 0.65 kcal/atom for the added atom, well above the 0.3 kcal/atom that is thought to be useful for drug optimization ^29,30^. Meanwhile, analogs that improved activity >10-fold had ligand efficiency of 1.4 for a single added atom (**Ext Data Figure 4**), close to the efficiency limit ^31^ (several even exceeded single atom ligand efficiencies of 2.0). Similarly, every analog that improved activity by ≥10-fold also improved lipophilic efficiency (**Ext. Data Figure 4**). More broadly, while there was little correlation between ΔclopgP and ΔpKi or ΔlogEC_50_ over the the 257 analogs (**Ext Data Figure 4b**), there was a strong linear correlation between both ligand efficiency and activity change and between lipophilic ligand efficiency and activity change—in both cases, substantially improved activity was accompanied by improved ligand and lipophilic efficiency. For analogs with much improved activity—especially those improved by >10-fold but also for those improved by >3-fold—the improvements cannot be simply laid at the door of increased hydrophobicity.

We investigated whether there were correlations between likelihood of potency improvement and ligand and binding site properties. One might expect that certain ligands or sites might better lend themselves to affinity/potency improvement, which would affect the domain of applicability of this study. For instance, weaker ligands might be easier to optimize, or smaller binding sites might be better suited to big affinity/potency jumps. We calculated if the parent molecular weight, binding site volume, the ligandability of the binding sites (calculated using Dscore^32^), or the affinity/potency of the starting parent compound was correlated with the likelihood of finding analogs with substantial affinity/potency improvement. Using the fraction of analogs achieving ≥10-fold improvement as the outcome, none of these properties showed a statistically meaningful correlation (**Figure 2b**). If one considers the magnitude of the affinity/potency change overall, and not just whether it improved ≥10-fold, correlations did emerge, with weaker vs stronger parents more likely to support improved affinity (R = −0.34, p = 3.3×10^−7^) and with larger binding sites also tending to do so (R = 0.23, p = 4.3×10^−4^) (parent size and site ligandability continued to show no association with affinity improvement). While these trends were echoed among matched pairs in ChEMBL (**Extended Data Figure 1**), the correlations overall were weak (even an R of -0.34 explains only 10% of set variance), and as activity changes drop below a 3-fold effect our confidence in the experimental values begins to drop. Taken together, these results suggest that: (1) as a strategy, systematic small perturbations can be unexpectedly successful at improving ligand affinity/potency^21,24^, perhaps even compared to modern design-heavy approaches^33^; (2) this effect is common across diverse parents and binding sites, (3) finds echo in the literature, and (4) random expectations for substantial affinity/potency improvement may be as high as 11% (95% CI 7.5%-15.2%).

To investigate the structural bases for the affinity/potency changes, we determined the structure of the parent inhibitors in complex with Mac1 and six analogs from the ’3122, 12 analogs from the ’3453 series, and four analogs from the ‘3794 series. These structures were determined to between 0.97 and 1.02 Å resolution (**Supplementary Table 2**), supporting atomic resolution analysis. The analogs with structures determined represented all six types of single-atom modifications in this study; 15 of them differed in affinity/potency from their parents by ≥3x, two by over 10-fold, and among themselves by up to 680-fold.

For some parents, analogs superposed well facilitating analysis of the effects. This was often true for the Mac1 ‘3453 (IC_50_ 8.9 µM) inhibitor series, for instance (**Figure 3a**). Displayed at the ortho-position of the phenyl ring the chloro of ‘0676 (IC_50_ 21 µM) projects into solvent and loses activity versus the parent (**Figure 3b**). Moving this chloro to the meta-position (’1304), where it packs with Phe132, Ile131, and Gly48, improves inhibition 11-fold (p-value <0.01) to 1.9 µM (**Figure 3c**). Moving this substitution one atom further over, to the para-position, had little effect (’6343, IC_50_ 2.1 µM) though a methyl at the same position (’9870, IC_50_ 7.6 µM) loses 4-fold affinity (p-value <0.01), presumably reflecting its poorer packing versus the chloro analog.

**Figure 3.**
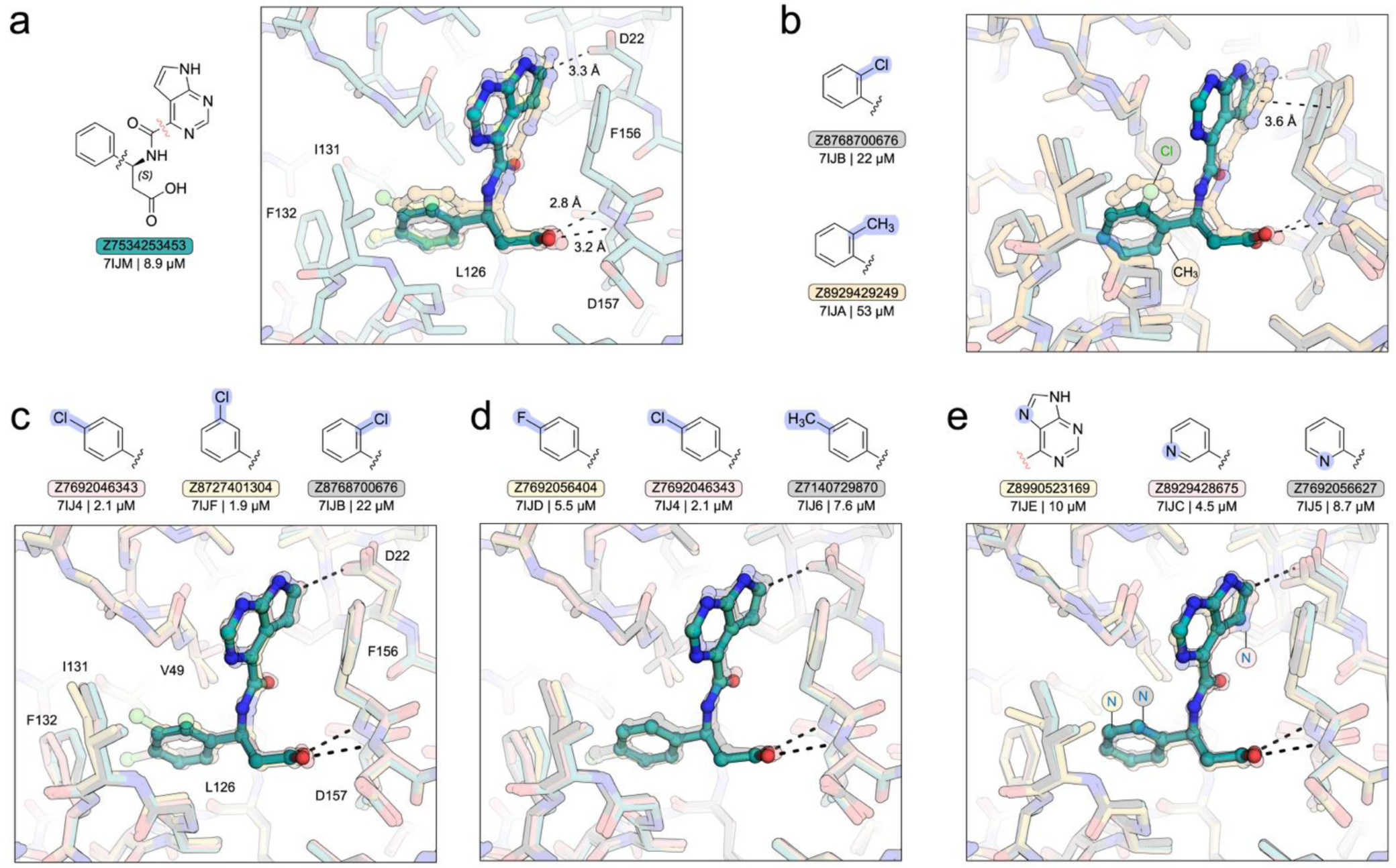
Structural basis of potency changes induced by single-atom modifications for Mac1 inhibitors. **a.** Chemical structure of the ’3453 parent and overlay of X-ray crystal structures of 12 analogs. **b.** Alignment of ‘0676 (21 μM) and ‘9249 (53 μM) with the ‘3453 parent showing the difference in binding pose for the ortho-chloro and ortho-methyl substituted analogs. **c.** Comparison of ‘6343 (2.1 μM), ‘1304 (1.9 μM) and ‘0676 (21 μM) reveals that repositioning the aryl chloride disrupts packing with I131 and F132, reducing potency. **d.** Para-methyl (‘9870, 7.6 μM), para-chloro (‘6343, 2.1 μM) substitutions have different effects despite similar sterics. Fluoro analog ‘6404 (5.5 μM) shows weaker binding than the ‘chloro (‘6343), likely reflecting weaker non-covalent interactions. **e.** Nitrogen substitutions (C→N) in ‘3169, ‘6627, and ‘8675 all pay increased desolvation penalties. Compared to their shared parent (8.8 μM), ‘8675 (meta-N) improved potency nearly 2-fold (4.5 μM), ‘6627 (8.7 μM) showed little improvement, and ‘3169 (10 μM) is slightly worsened.

If many of the effects of the small perturbation analogs could be explained *post hoc* from their structures, however, fewer were easily anticipated. Even the apparent simplicity of the ‘3453 series is belied by structural accommodations in the enzyme site. For instance, the para-chloro of the 2µM ‘6343 would have clashed with the enzyme without Phe132 and Ile131 rotating away. In doing so, Phe132 adopts a partially eclipsed conformation. Presumably the resulting strain is overcome by the improved packing of the buried para-chloro, but the outcome of this balance of terms seems difficult to intuit (encouragingly, free energy calculations did correctly predict that the para-and meta-chloro ‘6343 and ‘1304, respectively, would improve affinity while the ortho-chloro ‘0676 would lose it, though the agreement was more qualitative than quantitative; see below). Conversely, the methyl analog (‘9249, 53 µM) of ’0676 (IC_50_ 22 µM) is buried from solvent, not exposed, and might be expected to be more potent than its chloro cousin, but instead it loses another 2.5-fold in K_i_ (p-value <0.01; here, the opposite effect was anticipated by the free energy calculation, below). Meanwhile, in the complex between ‘4905 and the α2a adrenergic receptor, an added methyl group fits into a pre-existing sub-pocket and qualitatively its improved potency is reasonable. Quantitatively, its 52-fold effect on EC_50_ is at the outer edge of what might be expected by ligand-efficiency, especially since the methyl buries a serine hydroxyl. This is echoed by the free energy calculation, which here too correctly predicts improved activity, but only by 2- not 52-fold. These are examples of complexes where the analogs largely superpose on the parent and each other. In complexes where the single-atom changes led to substantial inhibitor movement, or to the adoption of multiple ligand conformations, prediction was harder still (**Extended Data Fig 5**). For example, the Mac1 methyl analog ‘3176 repositions the inhibitors in the pocket (**Extended Data Fig. 5b**), while compounds ‘9249 and ‘3194 adopt two distinct conformations in the site (**Extended Data Fig. 5b**), a phenomenon that is likely underreported in the PDB^34^ but is readily seen at ultra-high resolution. Taken together, this set of perturbations may provide an interesting and sometimes challenging test for prediction methods, and suggests that there may be merit in systematic small perturbations as a strategy, though showing that was not a goal of this study.

In optimizing chemical probes and leads, compound pharmacokinetics (PK) is as important as molecular target affinity/potency ^35,36^. We thus explored how the small perturbation analogs affected the PK properties versus those of the parent ligands, and how these changes related alterations of affinity/potency. We measured the following in vitro PK properties: metabolic stability in microsomes (half-life), plasma stability (half-life), plasma protein binding (fraction unbound), solubility (uM), hERG inhibition (IC50, uM), and membrane permeability (PAMPA assay) (cm/s) of both the parent ligands and their small change analogs (**Figure 4, Supplementary Table 3-8**). While these are all in vitro measurements, they are proxies that are considered predictive of in vivo PK^37–39^. Some of the small change analogs showed meaningful improvements relative to the parent ligands for each of these properties (**Extended Data Table 2**). For example, 12.4% of the analogs had ≥3-fold improvement in microsomal stability, while **20.8%** of them had a ≥3-fold improvement in plasma stability, versus the parents.

**Fig. 4.**
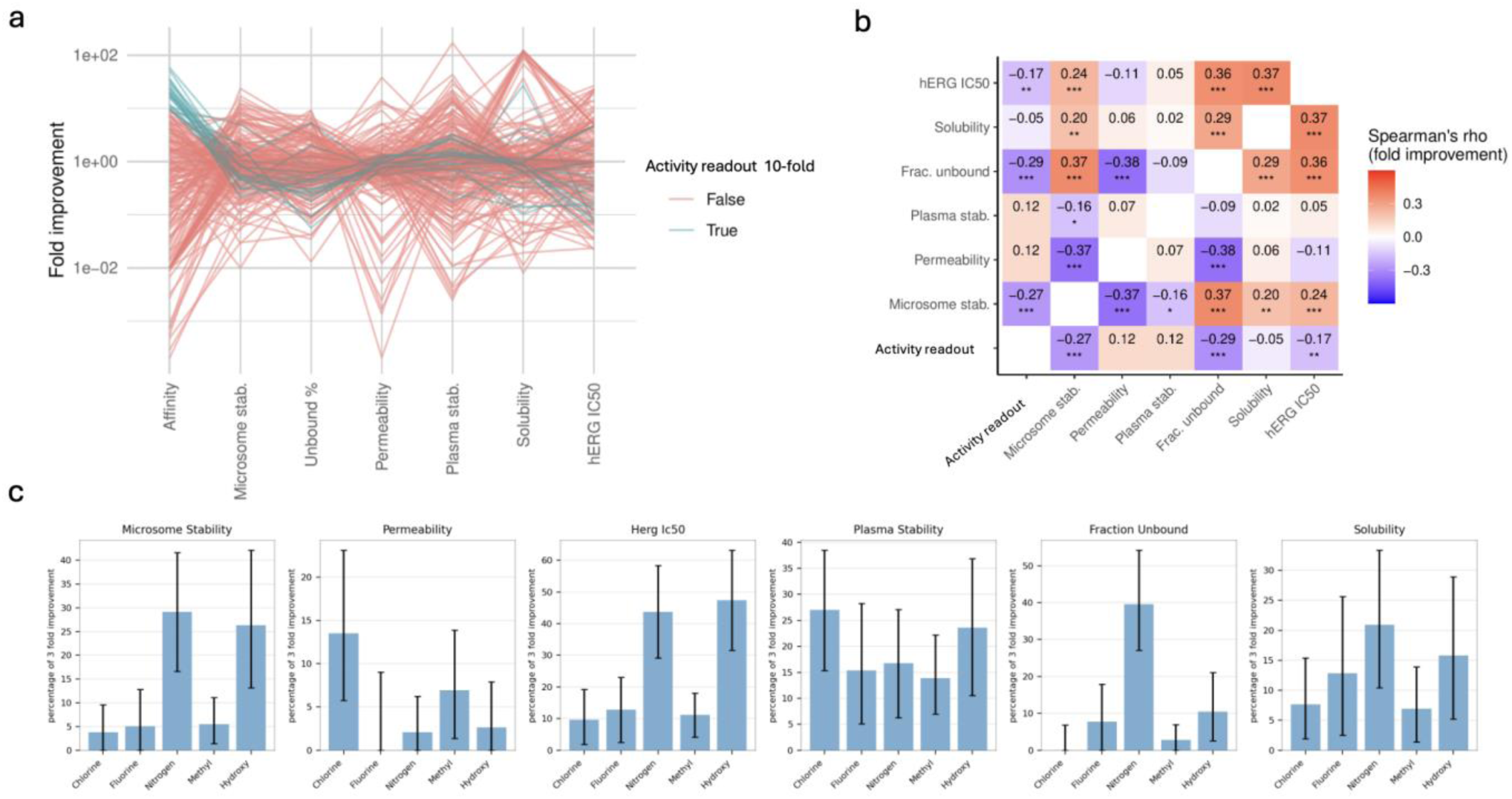
A frustrated landscape for ligand optimization. **a**, Fold-changes in pharmacokinetic properties for the 29 analogs that were ≥10-fold improved in affinity/potency versus a parent. **b**, Correlations between affinity/potency fold changes and those of the other PK properties. **c**, The fraction of single atom changes that were ≥3x fold improved in each parameter. Error bars represent 95% confidence intervals, estimated from 10,000 iterations of bootstrap resampling.

For **13.8%** of the analogs, solubility improved by ≥3-fold, while 6.2% of them improved ≥3-fold in permeability. Examining the fraction unbound in plasma, 13.2% of the analogs improved ≥3-fold. These findings illustrate that each property is amenable to meaningful optimization even by simple changes. We also observed a substantial fraction of analogs with ≥3-fold decreases in these same properties (**Extended Data Figure 6).** Considering what are perhaps the three most impactful in vitro PK measurements, microsomal stability, fraction unbound, and permeability, 36% of the analogs suffered >3-fold losses in at least one of these, almost all (96%) suffered at least some deterioration by the same criterion, and 37% deteriorated in all three in vitro PK properties.

If many of the small perturbation analogs improved substantially in individual PK properties, none of those with ≥10-fold increased potency improved or even maintained more than one of what we might consider the three most important ones: metabolic stability, PAMPA permeability, and fraction unbound in the plasma (**Figure 4.a, Extended Data Figure 7**); most were worse by all three properties. Indeed, the changes in in vitro PK were, at best, orthogonal to affinity/potency fold-change (**Figure 4b**), and several properties were anti-correlated with it.

This is born out on an atom-by-atom level: those atoms that most often improved affinity/potency substantially, like methyls and chloros (**Figure 2a**), were least associated with improved pharmacokinetic properties, such as microsomal stability or fraction unbound (**Figure 4c, Extended Data Figure 8**). Meanwhile, those atoms least likely to improve affinity/potency substantially, like aromatic C→N, were most likely to improve microsomal stability or fraction unbound. Density and cumulative distribution plots illustrate this divergence in effect (**Extended Data Figure 8**), with distinct shifts in property distributions across modifications. More broadly, most of the pharmacokinetic properties were largely independent of one another, with no correlations exceeding |ρ| > 0.5 and those that were statistically significant having Spearman ρ values ranging only from |0.16| to |0.38|. What emerges is a frustrated landscape for ligand optimization, with affinity/potency improvements counterbalanced by often worsening pharmacokinetics. While this trade-off has been noted in medicinal chemistry,^40,41^ this set quantifies its systematic impact.

The current benchmark only comprises several hundred analogs. If we could predict the results of the small perturbations, rather than having to experimentally test them, the set could be much extended. We therefore asked how well modern methods could predict the relative affinity/potency and pharmacokinetic changes we observed. In blinded experiments, we used free energy perturbation, arguably the highest level of theory available to the field, with the program FEP+ (Schrodinger, New York) to predict the changes in activity between the parents and analogs (**Figure 5 a-c**). Similarly, we asked how well we could predict in vitro pharmacokinetic changes between the parents and analogs, using widely accessible tools (**Figure 5d**).

**Figure 5:**
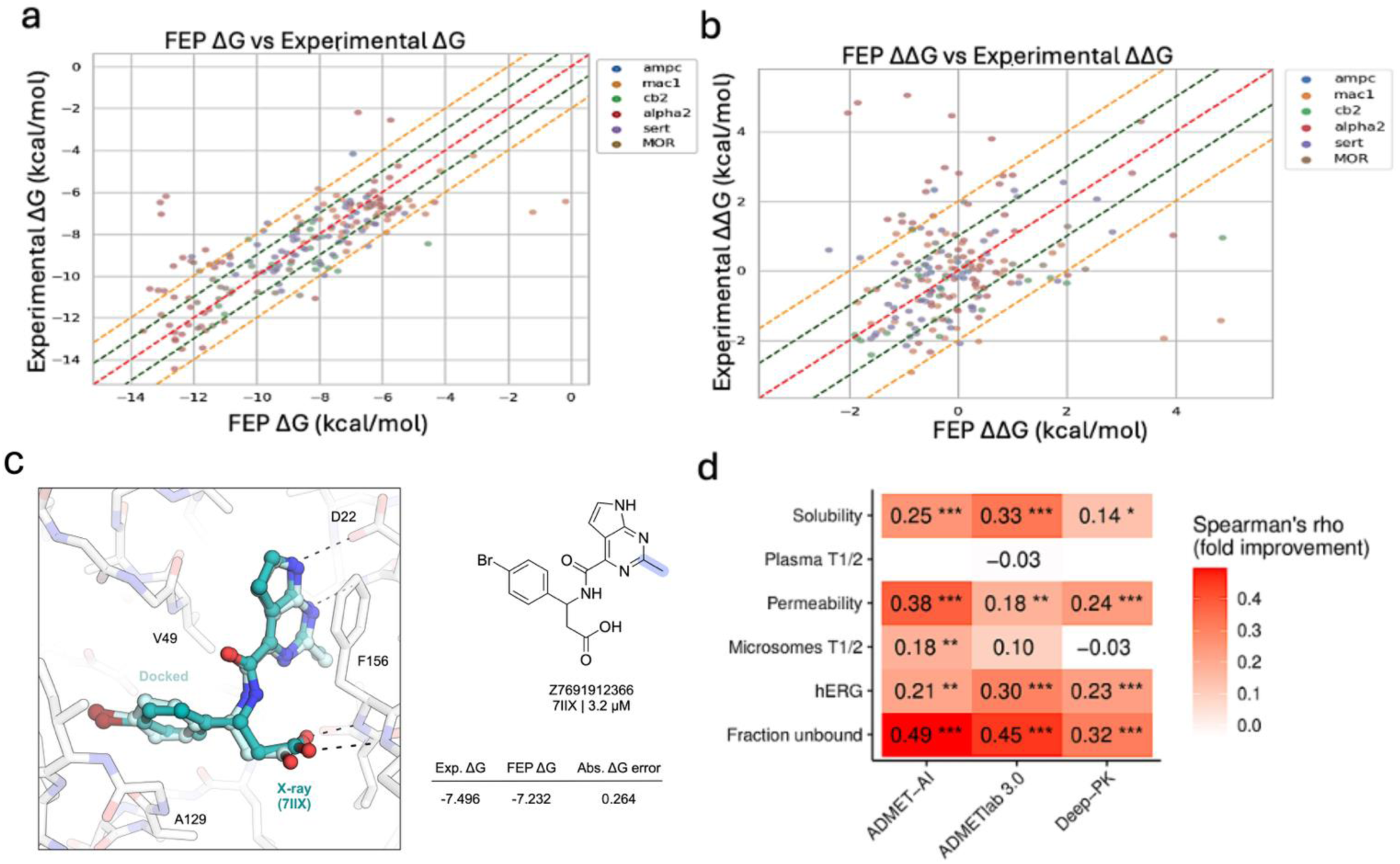
Accuracy of computational predictions of analog relative affinity and in vitro pharmacokinetics. **a**. Comparison of FEP-predicted and experimentally measured binding free energies (ΔG). Dashed lines indicate ±1 and ±2 kcal/mol error margins. **b**. Correlation between FEP-predicted and experimental measured relative binding free energy changes (ΔΔG). **c**. An example of good structural alignment between the predicted (cyan) and experimental (grey) poses, accompanied by a small error in the predicted ΔΔG. **d**. Heatmap of Spearman correlations between AI-predicted and experimentally measured fold changes across six in vitro PK properties, evaluated using three models. Color intensity indicates correlation strength; significance: p < 0.05 (*), < 0.01 (**), < 0.001 (***).

There was a strong overall correlation between the FEP+-predicted and experimentally measured binding free energies (ΔG), with most predictions (190) falling within ±2 kcal/mol of the experimental values, and 131 within ±1 kcal/mol, while the set had a mean unsigned error (MUE) to ΔG as 1.07 kcal/mol (**Figure 5a,b**). Of the 18 parents, the FEP+ predictions on 14 had ΔG MUE below 1.2 kcal/mol (**Supplementary Table 9**). If one considers the correlations qualitatively, of the 35 analogs predicted to lose affinity by 1 kcal/mol or more by FEP+ (i.e., in the range of confident prediction), 20 did lose affinity/potency by experiment. Meanwhile, of the 34 predicted to increase in affinity/potency by more than 1 kcal/mol by FEP+, 14 did so experimentally while 12 were experimentally neutral and 8 instead showed decreased potency. Categorization with a 1 kcal/mol threshold reveals a significant association between the direction and strength of FEP+ predictions and experimentally observed affinity/potency changes (χ² of 36.4 and a p-value < 0.001) (**Supplementary Table10**).

It is useful to note several of the challenges facing accurate FEP predictions for this series, which future work may share. The predictive power was likely weakened by our inability to measure affinity/potency for weak analogs, diminishing the overall range of affinities and making it impossible to measure affinities predicted to be very low by FEP+. We used an automated workflow to generate input poses for the calculations; on visual inspection, most of the outliers were caused by suboptimal initial poses of analogs. The accuracy of FEP calculations is strongly impacted by the quality of the ligand-complexed structures^42^, and the FEP+ predictions here began with docking poses, which are less reliable than experimental structures. For the 25 analog structures that we determined by crystallography (for Mac1 analogs) and by cryoEM (for an α2a analog, below), there were many cases where the predicted and experimental poses closely superimposed, and for these the FEP+ energies were largely consistent with the experimental measurements (Figure 5C).

Recently, AI-based platforms have been developed to facilitate ligand optimization by predicting ADMET properties with increasing accuracy and scale, guiding experiment^43,44^. We compared the ADMET predictions from ADMET-AI, ADMETLab 3.0, and Deep-PK^45–47^ to the experimentally measured values. The predictions of all three were significantly correlated with experimental measurements of plasma fraction unbound and with permeability, while microsomal stability, plasma stability and solubility, which are widely considered difficult to predict^48,49^, were essentially uncorrelated. Correlations in the 0.4 to 0.5 range, as they often were for fraction unbound and permeability, are strong enough to help guide compound selection in many circumstances, and these open-access tools may have broad impact. Still, even with a correlation of 0.53 in fraction unbound, the highest we observed, 28 analogs that had substantially increased fraction unbound experimentally were predicted to have *lower* levels than their parents, and 16 analogs that had higher protein binding than their parents experimentally were predicted to have less. Taken together, while the FEP and ADMET predictions can profitably model the effects of the small perturbation analogs, their correlations with experiment are not yet strong enough to allow this set to be extended by calculation alone.

### Multi-parameter optimization for an improved analgesic

Notwithstanding the frustrated landscape for ligand affinity/potency and pharmacokinetic improvement, medicinal chemists can often navigate the multi-parameter optimization to overall improve the properties of lead molecules. In this study, too, there were analogs that improved in affinity/potency with only modest sacrifice in pharmacokinetics (PK). Among these was an analog of an α2a parent, compound **‘4905**, where a single methyl group substitution improved potency 52-fold (**Figure 6a, b**). To understand this improvement at atomic resolution, we determined the structure of **‘4905**/alpha2a/G protein complex by single particle cryoEM to 2.8 Å resolution. Encouragingly, the experimental structure of **‘4905** superposed closely with the docking prediction for the parent ‘**3629** and with the experimental structure of a previously described lead member of this family of alpha2a agonists, ‘**9087,** suggesting that the methyl perturbation did not change the orientation of the analog, and that this family may be understood in the same structural context. From the cAMP dose–response curves, we observed a slight upward deflection at high concentrations for ‘4905. Consistent with this, PTX-treated cAMP assays showed a residual, concentration-dependent increase in cAMP for ‘4905 but not 3629 (**Supplementary Figure 6**), suggesting secondary Gs coupling by the ligand. Structurally, the extra methyl in 4905 fills a hydrophobic subpocket and shifts Y409^6.55^ (implicated in Gs activation), which may bias the receptor toward conformations permissive for Gs engagement. Improved steric complementarity with residues in TM5 and TM3, and a closer distance between the partly cationic bridging nitrogen with the crucial Asp128 on TM3, helps to explain the improved potency of **‘4905** over its parent.

**Figure 6.**
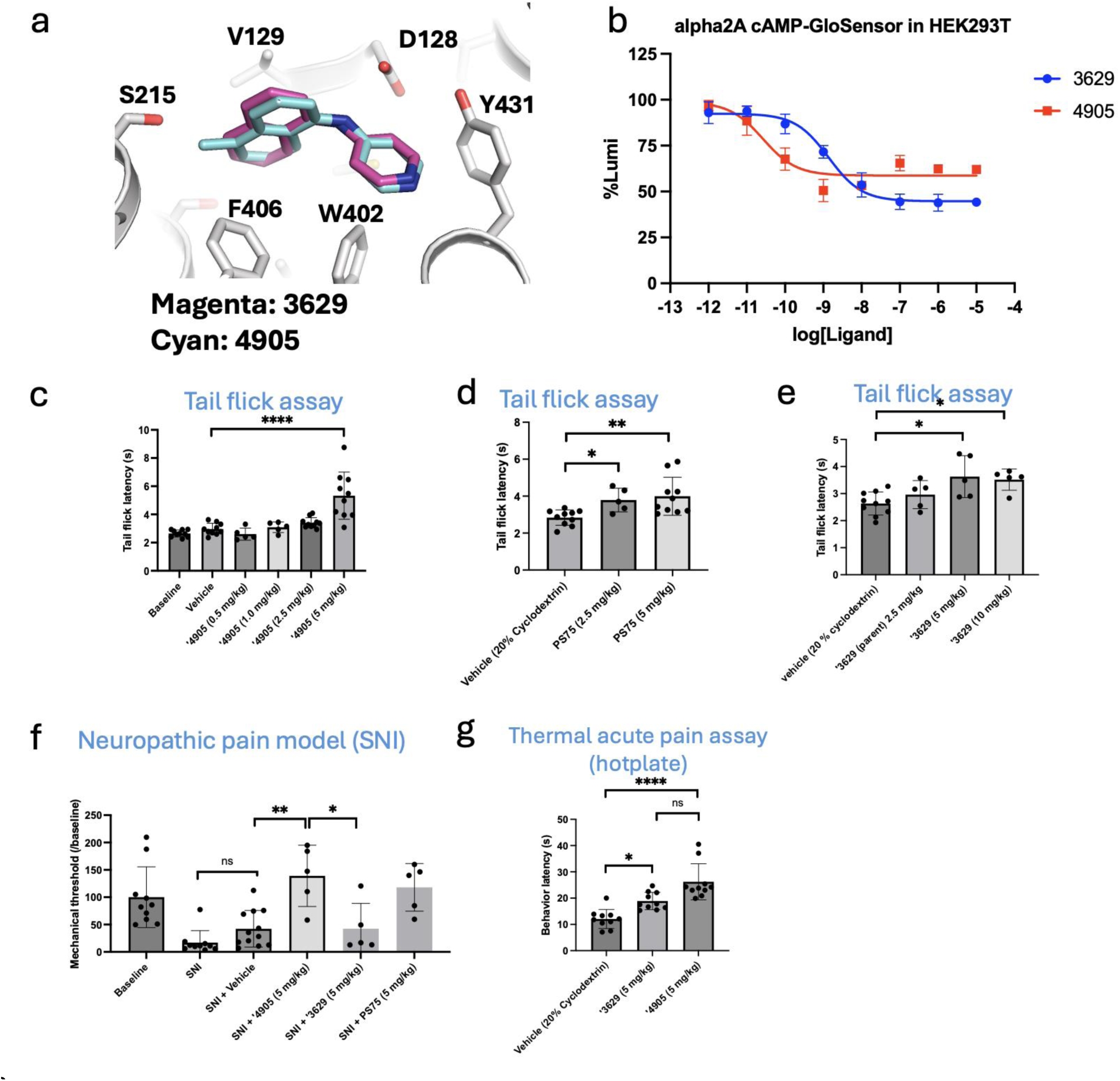
A single-atom addition to an alpha2a receptor agonist creates a more potent analgesic in vivo. **a.** Docking pose of parent ‘3629 superposed on the cryoEM structure of analog ‘4905, which differs from ‘3629 by the addition of a single methyl group. **b.** ‘4905 is 50-fold more potent than the parent ‘3629 in receptor activation in vitro. We note that the apparent reduced efficacy through Gi coupling at concentrations greater than 1 nM likely reflects the activation of Gs. **c**-**g**. ‘4905 is more potent than the parent ‘3629 in **c-e.** Reflex pain as measured in tail flick response to heat; **f.** Reversal of mechanical allodynia in a neuropathic pain model (SNI). **g.** Efficacy in reducing thermal acute pain assessed via the hotplate test.

While the improved potency of **‘4905** came at a cost to its in vitro pharmacokinetics, the effects were relatively modest: Compared to its parent, ‘**3629**, the fraction unbound decreased by 57% and the microsomal half-life was reduced by 41%. (**Supplementary Table 3-8**). The in vivo PK of **‘4905** qualitatively reflected the in vitro results, with CSF concentrations—a proxy for fraction unbound in the brain^50^—reduced by about 36%, while the half-life was reduced from 196 to 15.2 minutes (**Supplementary Table 3-8**). These size of these effects suggested that they might be outweighed by the analog’s great increase in activity. Accordingly, we compared the *in vivo* analgesia conferred by **‘4905** to its parent ‘**3629** and to **PS75**, the most potent analgesic known for this family of alpha2a agonists^51^. Despite the deterioration in pharmacokinetic properties, in mouse nociception assays including tail flick, neuropathic pain (SNI), and hot-plate the new analog ‘**4905** was far more potent than the parent **‘3629** and at least as potent as **PS75** (**Figure 6 c-e**). If ligand optimization is a frustrated landscape, it remains a navigable one as is well-known to medicinal chemistry^41^. In addition to beginning to provide a random background for such efforts, this study supports a strategy by which that landscape may be reconnoitered^21,24^.

## DISCUSSION

While ligand optimization is at the heart of drug discovery, quantifying the success of strategies to modify ligands has been difficult owing to a lack of a random background comparison set. Four key observations emerge from this study. **First,** systematic and unbiased small perturbations to 18 different scaffolds revealed that 11.2% of the analogs improved ≥10-fold in affinity/potency versus the parent (**Table 1**); “magic methyls”^25,52^ were common in this set. These improvements were observed against five of six receptors targeted, across physical properties of target and ligand, and across five orders of magnitude of parent affinity/potency. This begins to establish a background expectation for affinity/potency optimization against which methods and designs may be compared. It may also suggest a simple, unbiased approach for achieving substantial jumps in affinity/potency, improvements that will not always be obvious from structural analyses or even high-level simulations (**Figures 2** and **5**) but may complement them. **Second,** improved affinity/potency typically came at the cost of ligand pharmacokinetics (PK), with plasma fraction unbound, stability to liver microsomes, and permeability—among other terms—declining as analog affinity/potency increased. The individual in vitro pharmacokinetic terms were orthogonal to each other, and several were anti-correlated with affinity/potency improvement (**Figure 4**). This suggests a frustrated landscape for ligand optimization. Although the qualitative challenges and trade-offs in ligand optimization are known to medicinal chemists ^6,40,41^, this study begins to quantify them at scale and across a range of target classes and ligand properties. **Third**, computational prediction of affinity/potency and PK had some success anticipating the trends we observed (**Figure 5**) implying that they may eventually guide compound selection. Still, their correlations with experiment remained loose enough to preclude replacing systematic exploration and testing of molecules as this background set is further expanded, something seen with other potency prediction methods^53^. **Fourth,** while potency trades-off against PK, it is possible to find analogs where it rises sufficiently to overcome drops in exposure and half-life. A testament to this is the in vivo potency of the α2a analog **‘4905,** which despite suffering these trade-offs nevertheless is the most potent analgesic in vivo in a large series, and is certainly far more potent in vivo than its parent. It may be interesting to model the implications of the orthogonal and anti-correlated terms here to understand how best to navigate this multi-parameter optimization problem.

At first glance, this study is reminiscent of learnings from classical medicinal chemistry where conservative and ligand-based approaches like Topliss Trees^9^ dominated the field. But this would be to misremember the classic age of medicinal chemistry on two counts. First, the systematic small perturbations explored here were rarely used then, partly because the building blocks and reactions to support them were lacking. The ability to systematically explore small perturbations, across starting scaffolds and targets, reflects the advances in synthetic chemistry over the last 50 years and also the new building-block approaches that have expanded readily available compounds^54,55^. Second, the logic behind Topliss Trees and other quantitative structure-activity approaches^9,10^ were anchored in ligand physical chemistry without reference to receptor structure. For instance, in Topliss Trees the success of methyl and chloro derivatives were thought to teach opposite lessons because of their different electronic effects on the ligand, whereas the steric and non-polar properties of the two groups will seem similar from the view “inside the receptor”^52,56^. Indeed, this is what we find in the similar affinity/potency effects of the two substitutions. Meanwhile, the advantage of a random background, so integral to molecular biology, would have seemed foreign to classical medicinal chemistry. Thus, while a strategy of systematic small perturbations may smack of the “methyl, ethyl, propyl, butyl, futile” approach with which classic medicinal chemistry has been tarred ^57,58^, it has a different basis in theory and reflects new synthetic opportunities.

Certain limitations of this study merit airing. At 257 analogs and six targets, our set remains small and inevitably, if unintentionally, biased. We do not pretend that three class A GPCRs, one transporter, and two soluble enzymes adequately represent pharmacologically relevant targets, nor that the 18 parent ligands can adequately represent ligand space. Expanding the analysis to additional targets and ligand classes might reveal different trends. We note that the six targets do recognize a range of chemotypes, including large polar co-factors (Mac1), large hydrophobic lipids (CB2), anionic β-lactams (AmpC), cationic neurotransmitters (α2a and SERT), and peptides (µOR). Experimentally, we assessed activity changes using radioligand competition binding assays (SERT, α2a), second messenger assays (α2a, CB2, and MOR), and binding and enzyme activity (Mac1 and AmpC) assays. While ligand competition correlates with K_i_, agonist EC_50_s are affected by receptor expression. Because we compared the relative activities of analogs versus their parents, controlled for receptor expression (**Supplementary Figure 4**), and measured effects in a low expression level domain (**Supplementary Figure 5**), the effects of receptor expression on relative activity should be modest, though they cannot be completely discounted. For these and related reasons, changes of EC_50_ between agonist parent and analogs cannot be read as changes in affinity the way that changes in Ki can be, though EC_50_ changes remain the relevant metric for agonist activity. Overall, the impact of these effects may be inspected case by case in full concentration-response for any parent ligand pair with >3-fold improvement in activity (**Supplementary Figure 2**). In the area of in vitro pharmacokinetics, our use of liver microsomes rather than hepatocytes to measure metabolic stability meant that some types of metabolism were missed, including glucuronidation. This might make groups like phenolic hydroxyls seem less labile than they would be in vivo. More broadly, for all but two molecules we only measured in vitro not in vivo pharmacokinetics. Whereas the in vitro measurements qualitatively anticipated the in vivo effects where we did measure them, and are widely used in ligand optimization, their quantitative prediction of in vivo behavior is only approximate.

These limitations should not obscure the main observations of this study. Over 11.2% of systematic and unbiased small perturbations improved analog affinity/potency ≥10-fold, beginning to establish a background expectation for the likelihood of substantial ligand affinity/potency improvement and a systematic approach to doing so ^21,24^. Balancing this was a concomitant deterioration in ligand pharmacokinetics, which will lower the exposure and half-life of a ligand in vivo, counteracting the improvements in affinity/potency. Navigating this multi-parameter space is at the heart of medicinal chemistry; this study supports to the development of quantitative models to do so.

## Methods

### ChEMBL database molecular pairs

From the PostgreSQL version of CHEMBL34^23^ (https://doi.org/10.6019/CHEMBL.database.34), we filtered the database for compounds with activities reported against a single protein, with either Ki, Kd, IC_50_ or EC_50_ as the activity type. To be considered a molecular pair, compounds had to have the same assay ID, the same target ID, the same activity type, and the same reference document (publication) ID, in addition to having a parent-analog relationship as defined in this work, meaning a C-H group replaced by C-OH, C-F, C-Cl, C-Br, C-CH3 or an aromatic carbon replaced by a nitrogen. This left us with 191,732 parent-analog pairs.

### Analog enumeration and synthesis

Eighteen parent compounds were selected from previously published literature^51,59,60^ or datasets (https://asapdiscovery.org/outputs/molecules/#ASAP-SARS-COV-2-NSP3-MAC1) based on their known binding activity, structural relevance, or representation of diverse chemical scaffolds. Starting from a parent compound, we used RDKit (www.rdkit.org) to identify all C-H bonds and iteratively replace the hydrogen atom with a methyl, hydroxyl, fluoro, chloro or bromo group. We also identified aromatic carbons with two heavy-atom neighbors and replaced them with nitrogen. Every analog generated was represented as a canonical isomeric SMILES string and added to a set to remove duplicates. For each of the 18 parents, the full set of possible analogs that could be synthesized for < $400 for 10mg were ordered.

### Confidence intervals and statistics

When reported, 95% confidence intervals were derived from bootstrap resampling with 10,000 iterations. In cases where the observed frequency of success is exactly 0, bootstrap will fail to give an upper bound for the interval. In such cases, we used the Clopper-Pearson method to estimate an upper bound based on the sample size.^61^ Pearson and Spearman correlations with their associated p-values were computed using the scipy.stats module from the SciPy package^62^.

### FEP simulations

These were conducted using FEP+ within the Schrödinger software suite (versions 2025-2) with the OPLS4 force field^63^ and the modified SPC water model. The default setting was used for the number of lambda windows selection where it depends on the type of perturbations; charge-changing, core hopping, and all other perturbations have 24, 16, and 12 lambda windows respectively. For alchemical transformations with charge changes, the total charge of the simulation box was kept constant by transmuting a Na+ or Cl- ion to water or vice versa. In addition, a 0.15 M concentration of NaCl was added to the simulation box of charge-changing perturbations. For α2A, CB2, and SERT, the FEP+ membrane protocol was applied where a POPC membrane was added to the system in simulation. All other settings were kept default except the simulation time was extended from 5 ns to 10 ns.

The default FEP map generation protocol was used with the parent compound selected as the biased node. In preparing proteins and ligands for FEP+, the Schrödinger protein preparation workflow and LigPrep were used. The initial binding poses of parent compounds were from poses generated by DOCK3.8, and analogs were aligned to the parents with severe steric clashes resolved using the FEP+ Pose Builder workflow.

### Docking

Whereas no structural information was used in the design of analogs from parent compounds, for FEP+ calculations and for post hoc structural analysis, we generated ligand bound complexes of the parents and relevant ligands using DOCK.3.8^54,64^. Ligands were docked into the receptor binding site using grids prepared in previous studies^51,59,60,65,66^.

### Assay selection

We use one consistent readout per parent series (no mixing within a series). SERT and AmpC are reported as Ki; Mac1 is reported as IC50 from peptide displacement (a scalable functional hydrolysis assay is unavailable); GPCR agonist series are reported as EC50 and antagonist series as Ki, such that MOR is EC50-only whereas α2A and CB2 include EC50 and Ki depending on the parent series. Representative concentration–response curves and the corresponding Z values for each assay are provided in **Supplementary Figure 3**.

### Transporter assays for SERT Ki

SERT activity was measured using the Neurotransmitter Transporter Uptake Assay Kit from Molecular Devices (Catalog #R8174), following the manufacturer’s protocol with slight modifications as described previously^65^.

HEK293 cells stably expressing human SERT were plated in poly-L-lysine (PLL)-coated 384-well black, clear-bottom plates at a density of 15,000 cells in 40 µL per well, using DMEM supplemented with 1% dialyzed FBS (dFBS). Cells were incubated overnight at 37°C with 5% CO₂ to allow adherence and recovery. The following day, the medium was carefully aspirated, and cells were incubated with 25 µL per well of test compound solutions prepared in assay buffer (1× HBSS, 20 mM HEPES, pH 7.4, supplemented with 1 mg/mL BSA) for 30 minutes at 37°C. After drug treatment, 25 µL per well of dye solution (as provided in the kit) was added directly to the wells, followed by an additional 30-minute incubation at 37°C. Fluoxetine (10 µM) was used as a positive control for SERT inhibition. Fluorescence was measured using the FlexStation II microplate reader with excitation at 440 nm and emission at 520 nm.

Relative fluorescence units (RFUs) were exported and analyzed using GraphPad Prism 10.0 to calculate IC₅₀ values, from which Ki values were derived using the Cheng-Prusoff equation.

### CB2 radioligand binding assay

CB2 receptor binding assays were performed using membrane preparations from HEK293 cells stably expressing human CB2, following previously published methods^67,68^. Membranes were resuspended in TME buffer containing 0.1% BSA (w/v) and 25 µg of membrane protein was added per well. The assay was conducted using the radioligand [³H]CP-55,940 at a final concentration of 0.75 nM, prepared in assay buffer. Nonspecific binding was defined in the presence of 5 µM unlabeled CP-55,940. Test compounds were applied at increasing concentrations to assess competition.

Reactions were incubated at 30 °C for 1 hour with gentle shaking. After incubation, samples were transferred to Unifilter GF/B 96-well filter plates and filtered using a Packard Filtermate-196 cell harvester (PerkinElmer). Plates were washed four times with ice-cold wash buffer (50 mM Tris-HCl, 5 mM MgCl₂, 0.5% BSA, pH 7.4). Radioactivity bound to the filters was quantified via liquid scintillation counting. Specific binding was calculated by subtracting nonspecific from total binding. IC₅₀ and Ki values were calculated using nonlinear regression in GraphPad Prism 9 using the Cheng-Prusoff equation.

### α2A Receptor Binding Assay

α2A adrenergic receptor binding was performed using membrane preparations from insect cells expressing human α2A receptors, as previously described^51^. Membranes were incubated with increasing concentrations of test compounds and 5 nM [³H]-rauwolscine in buffer containing 20 mM HEPES (pH 7.5) and 100 mM NaCl, at room temperature for 2 hours. After incubation, samples were filtered onto GF/B filter plates, washed with ice-cold buffer, and radioactivity was quantified by liquid scintillation counting. IC₅₀ and Ki values were derived using nonlinear regression in GraphPad Prism.

### GloSensor cAMP assay for α2A, CB2, and MOR

The GloSensor cAMP assay was performed following the manufacturer’s instructions (Promega) with slight modifications as previously described^51,69^. Briefly, wild-type (WT) human α2A, CB2, and MOR were cloned into the pcDNA3.1 vector and co-transfected with the 22F cAMP GloSensor plasmid into HEK293T cells cultured in 6-well plates. After 24 hours, cells were reseeded into 96-well white plates in CO₂-independent medium and equilibrated with GloSensor cAMP reagent as per the manufacturer’s protocol. Cells were incubated for 1 hour at 37 °C followed by 1 hour at room temperature. Where applicable, 10 μM forskolin was used to elevate basal cAMP levels for assessing receptor-mediated inhibition. Serially diluted test compounds were added, and luminescence signals were recorded using a PerkinElmer microplate reader. Data were analyzed using GraphPad Prism 9.0 to calculate EC₅₀ or IC₅₀ values.

### AmpC β-lactamase Inhibition Assay

The AmpC β-lactamase inhibition assay was performed as previously described^60^. The candidate inhibitors were dissolved in DMSO (20 mM stock) and diluted to maintain a constant 1% DMSO (v/v) in 50 mM sodium cacodylate buffer (pH 6.5).

Assays were performed in the presence of 0.01% Triton X-100 to reduce aggregation artifacts. AmpC enzymatic activity was monitored spectrophotometrically using CENTA or nitrocefin as substrates.

Initial screening was performed at 200 µM, 100 µM, and 40 µM compound concentrations. Substrate concentrations were selected based on known Km values to achieve defined [S]/Km ratios: for CENTA ([S] = 50 µM, Km = 27.6 µM) and nitrocefin ([S] = 100 µM or 28 µM, Km = 180 µM). Reactions were carried out in 96-well format on a BMG Labtech CLARIOstar plate reader, with substrate and enzyme injected into wells containing the inhibitor, followed by kinetic measurement over 50 seconds. IC₅₀ values were determined by fitting inhibition curves in GraphPad Prism using a fixed Hill coefficient of 1, and Ki values were calculated using the Cheng-Prusoff equation^70^.

### HTRF assay for Mac1

Binding of the compounds to Mac1 was assessed by the displacement of an ADPr conjugated biotin peptide from His6-tagged protein using a HTRF-based assay, as previously described^71^.The expression sequences used for SARS-CoV-2 Mac1 are listed below. All proteins were expressed and purified as described previously for SARS-CoV-2 Mac1^71^. Compounds were dispensed into ProxiPlate-384 Plus (PerkinElmer) assay plates using an Echo 650 Liquid Handler (Beckman Coulter). Binding assays were conducted in a final volume of 16 μl with 12.5 nM NSP3 Mac1 protein, 200 nM peptide ARTK(Bio)QTARK(Aoa- RADP)S (Cambridge Peptides), 1:20000 Anti-His6-Eu3+ cryptate (HTRF donor, PerkinElmer AD0402) and 1:500 Streptavidin-XL665 (HTRF acceptor, PerkinElmer 610SAXLB) in assay buffer (25 mM 4-(2-hydroxyethyl)-1-piperazi-neethanesulfonic acid (HEPES) pH 7.0, 20 mM NaCl, 0.05% bovine serum albumin and 0.05% Tween-20, the latter also to reduce aggregation artifacts). Assay reagents were dispensed manually into plates using an electronic multichannel pipette. Mac1 and peptide were preincubated for 30 min at room temperature before HTRF reagents were added.

Fluorescence was measured after a 1 hour incubation at room temperature using a Perkin Elmer EnVision 2105-0010 Dual Detector Multimode microplate reader with dual emission protocol (A = excitation of 320 nm, emission of 665 nm, and B = excitation of 320 nm, emission of 620 nm). Compounds were tested in triplicate in a 14-point dose response. Raw data were processed to give an HTRF ratio (channel A/B × 10,000), which was used to generate IC_50_ curves using nonlinear regression using GraphPad Prism v.10.0.2 (GraphPad Software, CA, USA).

### hERG channel inhibition

hERG channel inhibition was evaluated using a Thallium Flux assay on HEK293 cells stably expressing the human Ether-à-go-go Related Gene (hERG) potassium channel. Cells were seeded at a density of 8000 cells per well in 384-well poly-D-lysine–coated plates and incubated for 24 hours under standard conditions (37 °C, 5% CO₂). The next day, a thallium-sensitive dye was added to the cells, followed by a 1-hour incubation to ensure dye uptake. Test compounds were added to achieve a final concentration of 30 μM in 0.5% DMSO, and the cells were incubated for an additional 30 minutes at room temperature. Subsequently, a stimulation buffer containing thallium was added and fluorescence measurements were taken using a FLIPR Tetra system. Data were collected every 3 seconds for 3 minutes (excitation: 470–495 nm, emission: 515–575 nm).

Fluorescence intensity over time was analyzed to calculate the area under the curve (AUC) from which percentage inhibition was determined versus haloperidol at 100μM (positive control) and vehicle (DMSO).

### α2A Receptor Purification and Structure Determination

Wild-type human α2A adrenergic receptor (α_2A_AR) was cloned into a pVL1392 vector with an N-terminal FLAG tag. The construct was expressed in *Spodoptera frugiperda* (Sf9) insect cells using the BestBac system. Cells at a density of 4 × 10⁶ cells/mL were infected with virus and incubated for 48 hours at 27 °C. The receptor was solubilized and purified by FLAG affinity chromatography and size-exclusion chromatography in the presence of 10 μM compound ‘4905. Monomeric peak fractions were concentrated used for G protein complex formation. G_αo_G_β1γ2_ heterotrimeric G proteins were expressed in Hi5 insect cells and purified using Ni²⁺-affinity following detergent solubilization and dephosphorylation. The final α_2A_AR–G_αo_G_β1γ2_–scFv16^72^ complex was assembled in the presence of ‘4905 and purified by size-exclusion chromatography. Cryo-EM grids were prepared using UltrAufoil R1.2/1.3 300 mesh grids and vitrified in liquid ethane. Data were collected on a Titan Krios G3 electron microscope equipped with a K3 direct electron detector. Image processing was performed using cryoSPARC, yielding a final reconstruction at ∼2.8 Å resolution. Model building and refinement was performed using PDB 7EJ8 as a starting model using ChimeraX^73^, Phenix^74^, and Coot^75^. The final structures have been deposited in the PDB with accession codes 9PLO and 9PLN.

### Mac1 Purification and Crystallization

Wild-type Mac1 protein (P4_3_ construct, residues 3–169) was expressed in *E. coli* BL21(DE3) as an N-terminal His₆-tagged construct and purified by Ni²⁺ affinity chromatography ^76^. The His-tag was cleaved with TEV protease, followed by size-exclusion chromatography (Superdex 75) in 20 mM Tris-HCl (pH 7.5), 150 mM NaCl, and 1 mM DTT. Purified protein was concentrated to 40 mg/ml for crystallization and stored at -80°C.

Crystals were obtained by sitting-drop vapor diffusion in 28% PEG 3000 and 100 mM CHES (pH 9.5). Compounds (100 mM in DMSO) were added to crystal drops using an Echo 650 acoustic dispenser to a final concentration of 10 mM. Crystals were incubated at room temperature for 2–4 hours and vitrified in liquid nitrogen without additional cryoprotection. X-ray diffraction data were collected at ALS (beamline 8.3.1) and processed using XDS and Aimless. Structures were determined to resolutions ranging from 0.97 to 1.02 Å. Ligands with low occupancy or conformational disorder were modeled using PanDDA and refined using Phenix ^77^.PanDDA event maps are shown in Supplementary Figure 7. The structure of ‘3184 was determined using a racemic preparation of the compound (‘9037). X-ray data collection and refinement statistics are summarized in **SI Table 2**, as are the 32 PDB IDs.

### Microsomal Stability

Microsomal stability of compounds was evaluated using pooled mouse liver microsomes (XenoTech, M3000/lot #2010026) to estimate their metabolic stability and predict hepatic clearance. Each compound was incubated at 2 μM in a reaction mixture containing 0.42 mg/mL microsomal protein, phosphate buffer (100 mM, pH 7.4), MgCl₂ (3.3 mM), NADPH (3 mM), glucose-6-phosphate (5.3 mM), and glucose-6-phosphate dehydrogenase (0.67 units/mL). Reactions were conducted at 37 °C in 96-well plates with shaking at 100 rpm. Samples were collected at five time points (0, 7, 15, 25, and 40 minutes), and reactions were quenched by adding five volumes of acetonitrile containing an internal standard. After centrifugation at 5500 rpm for 5 minutes, the supernatants were analyzed via HPLC-MS/MS. The elimination rate constant (kel), half-life (t₁/₂), and intrinsic clearance (Clint) were calculated by plotting the natural logarithm of the remaining parent compound versus time. Stability was compared with reference standards such as imipramine and propranolol.

### Plasma Protein Binding (PPB)

Plasma protein binding (PPB) was measured using equilibrium dialysis with a 14 kDa molecular weight cut-off membrane in a 96-well HTD96b dialyzer. Mouse plasma containing 1 μM test compound (0.005% DMSO, 1% acetonitrile) was placed in one chamber, and phosphate-buffered saline (PBS, pH 7.4) in the opposing chamber. The assembled plates were incubated at 37 °C with 5% CO₂ and ∼95% humidity, shaking at 250 rpm for 5 hours to reach equilibrium. After incubation, aliquots from each chamber were mixed with equal volumes of the blank opposite matrix and processed with acetonitrile containing internal standard. Supernatants obtained after centrifugation were analyzed by HPLC-MS/MS. The percentage of compound bound to plasma proteins was calculated using the peak area ratio in buffer to plasma compartments. Recovery and stability standards were included to ensure accuracy and reliability. Verapamil served as a reference control. Most compounds showed moderate binding, typically ranging between 75–85%.

### Plasma stability

Plasma stability was assessed in non-sterile mouse plasma (Li-heparin treated) at 1 μM concentration (final DMSO content was 0.005%). Incubations were carried out in aliquots of 60 μL each (two per time point) at 37 °C under 5% CO₂ and high humidity (∼95%). The reactions were quenched with 240 µL of 90% acetonitrile containing an internal standard at 0, 20, 40, 60, and 120 minutes, followed by centrifugation at 6000 rpm for 5 minutes. Supernatants were analyzed via HPLC-MS/MS to determine the percentage of parent compound remaining at each time point. Data were plotted to calculate compound half-lives (t₁/₂). Reference compounds, verapamil and propantheline, were used as high and low stability controls, respectively. This assay is critical for identifying compounds susceptible to degradation by plasma esterases or hydrolytic enzymes, helping inform pharmacokinetic optimization strategies during lead selection.

### Thermodynamic Solubility

Aqueous thermodynamic solubility was determined in PBS (pH 7.4) for 247 compounds using a shake-flask method followed by UV absorbance quantification. Dry powder compounds were dissolved in PBS to a theoretical concentration of 4 mM and incubated in duplicates at 25 °C for 4 and 24 hours with shaking. After incubation, samples were filtered using HTS 96-well filter plates. The filtrates were diluted 2-fold in acetonitrile with 4% DMSO for UV analysis. The incubation samples for charged molecules were additionally diluted 10-fold with 50% acetonitrile/PBS with 2% final DMSO. Calibration curves (0–200 μM) were prepared in 50% acetonitrile/PBS (2% final DMSO). Absorbance was measured between 230–550 nm using a SpectraMax Plus microplate reader. Compound-specific absorbance maxima were used to calculate concentration using SoftMax Pro and Excel. The assay dynamic range is ∼2–400 μM (∼20–4000 μM for charged molecules), with values near the upper limit treated as semi-quantitative. Ondansetron was used as a reference compound. This method reflects equilibrium solubility under physiologically relevant conditions and helps rank compounds for formulation feasibility.

### PAMPA-BBB

Passive blood-brain barrier (BBB) permeability was estimated using a Parallel Artificial Membrane Permeability Assay (PAMPA-BBB) with a phospholipid-coated membrane simulating the brain endothelium. Test compounds (50 μM in Prisma HT buffer, pH 7.4, with 0.5% DMSO) were added to donor wells, while brain sink buffer was added to the acceptor wells. The donor and acceptor chambers were separated by a 0.45 µm filter membrane coated with brain polar lipids. Plates were incubated without agitation at room temperature for 4 hours. Post-incubation, samples from both chambers, as well as a standard solution, were diluted with acetonitrile containing an internal standard. Apparent permeability coefficients (log Papp) were calculated based on peak area ratio. Clozapine and chlorpromazine were used as high-permeability controls, while ranitidine represented low permeability. The assay provides a high-throughput, non-cell-based method for estimating CNS exposure potential.

### Behavioral analyses

All animal behavior experiments were conducted with the experimenter blinded to treatment. Mice were habituated individually in Plexiglas enclosures for 1 hour prior to testing. Compounds were administered subcutaneously 30 minutes before behavioral assessment, and where applicable the α2A adrenergic receptor antagonist atipamezole (2 mg/kg, intraperitoneally) was given 15 minutes before compound injection. Tail flick latency was measured by immersing the distal third of the tail in a 50°C water bath and recording the withdrawal time. For the neuropathic pain model, spared nerve injury (SNI) was performed under isoflurane anesthesia by ligating and transecting two of the three branches of the sciatic nerve, sparing the sural nerve. Mechanical thresholds were assessed 7 to 14 days post-surgery using von Frey filaments and the up-down method, and values were normalized to each animal’s baseline. Thermal nociception was evaluated using a 55°C hot plate, and the latency to nocifensive behavior (paw lick or jump) was recorded with a cutoff of 45 seconds to prevent tissue damage. Behavioral data are summarized as group size (n), mean, and SD (**Supplementary Table 12**). For the SNI experiment, we used two-way ANOVA followed by Tukey’s post-hoc multiple comparisons. For the hotplate assay, we used one-way repeated-measures ANOVA with Friedman post-hoc testing. For tail-flick, the 4905 dose–response was analyzed by two-way ANOVA with Dunnett’s post-hoc comparisons to vehicle/control, whereas PS75 and 3629 dose groups were analyzed by one-way ANOVA with Kruskal–Wallis post-hoc testing (**Supplementary Table 12**).For sample size, we did not perform formal a priori power calculations. Instead, group sizes were guided by our prior experience with these assays and by precedent in the literature. This approach may limit sensitivity to small effects. Raw data for the animal assays are provided in **Supplementary Table 13**.

## Data availability

Most primary data is available through this manuscript and its extended and supplementary materials. Crystal and cryoEM structures and supporting electron density are being made available via the Protein Data Bank, including PDB 9PLN, the structure of the α2a/’4905 complex and PDB 14AB, 7IIW, 7IIX, 7IIY, 7IIZ, 7IJ0, 7IJ1, 7IJ2, 7IJ3, 7IJ4, 7IJ5, 7IJ6, 7IJ7, 7IJ8, 7IJ9, 7IJA, 7IJB, 7IJC, 7IJD, 7IJE, 7IJF, 7IJG, 7IJH, 7IJI, 7IJJ, 7IJK, 7IJL, 14AM, 14AN, 14AO, 14AP, and 7IJM, structures of Mac1 in complex with inhibitors..

## Code availability

ZINC tools used in the selection of the small perturbation analogs are openly available and zinc22.docking.org.

## Acknowledgements

Supported by US DARPA grant HR0011-19-2-0020 (to B.K.S., A.M., A.I.B., and B.L.R.), by US NIH R35GM122481 (to B.K.S.) by US ARPA-H grant 1AY1AX000035 (PI J.S.F.), O.M. partially supported by US NIH postdoctoral fellowship F32GM154469. We thank Yeyue Xiong for help with the FEP studies. We thank the UCSF Cryo-EM facility staff for training and technical assistance. UCSF Cryo-EM equipment is partially supported by NIH grants S10OD020054, S10OD021741, and S10OD026881, and by the Howard Hughes Medical Institute.

## Author contributions

B.K.S., X.X., and O.M. designed the project. X.X. and O.M. designed the analogs with help from N.L. Alpha₂ receptor studies were performed by X.X. and H.H., with guidance from P.G. Alpha2A-related behavioral analyses were conducted by J.B., with guidance from A.I.B. Alpha₂ structural studies were performed by K.S., under the supervision of A.M. SERT binding studies were carried out by X.P.H. and J.W., with guidance from B.L.R. and ligand choice from Y.W. Mac1 biochemical assays were conducted by Y.U.D. M.S and M.D. supervised by A.A., and Mac1 crystallography was performed by G.C. with guidance from J.S.F. CB2 receptor assays were performed by X.X. and C.T., with ligand advice from M.R. AmpC assays were conducted by X.X. and F.L. All in vitro ADME and safety assays were supported by Y.H. and Y.K. through Bienta. FEP calculations and analysis were performed by DS guided by GZ and RA. O.M. performed statistical analyses for both ChEMBL and experimental data. X.X., O.M., and B.K.S. prepared the manuscript. B.K.S. supervised the project. All authors reviewed and approved the final manuscript.

---

**Extended Data Fig. 1.**
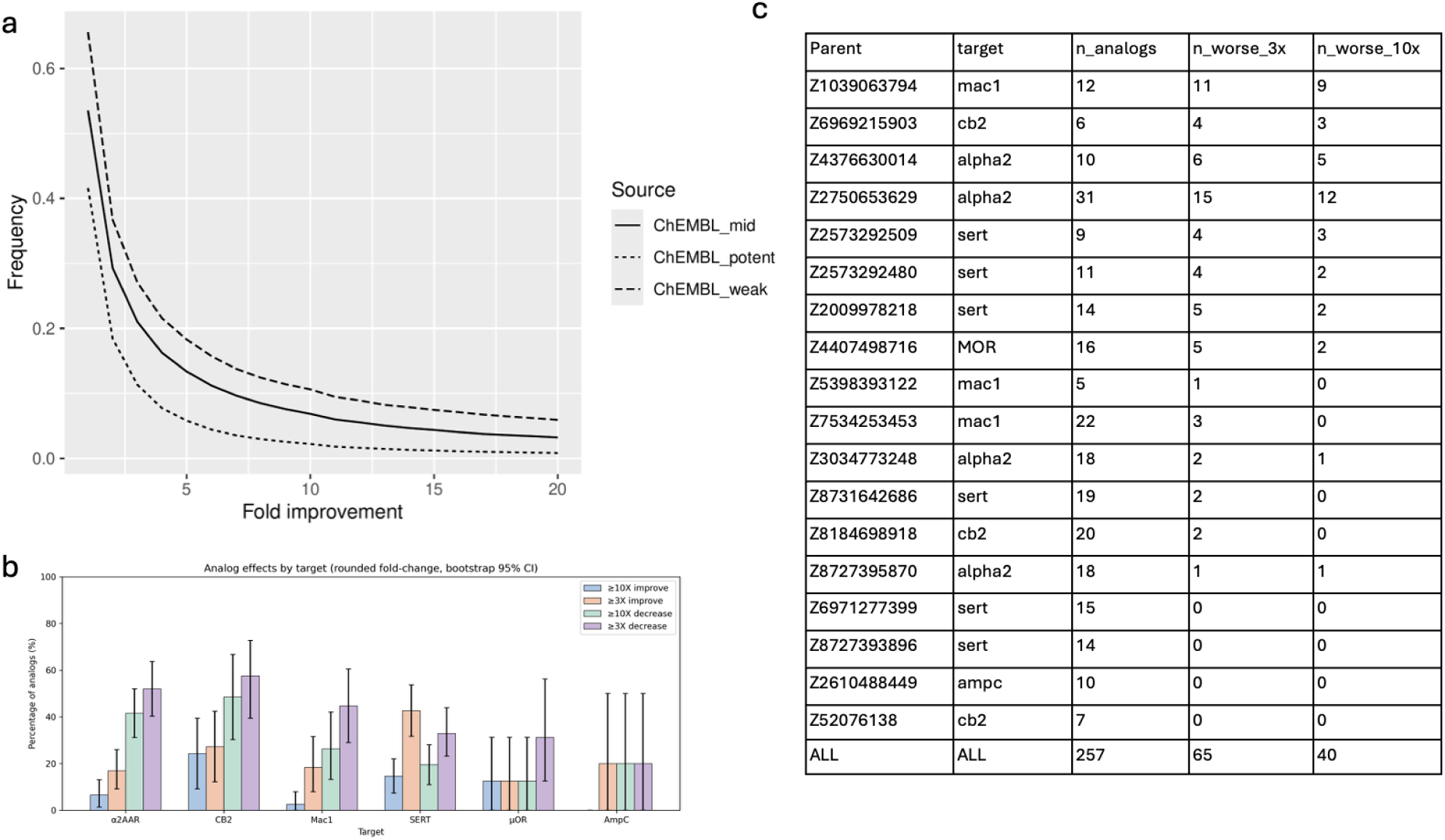
Frequency of activity changes from single-atom substitutions in ChEMBL and in this study. **a,** Cumulative frequency of single non-hydrogen atom substitutions (CH_3_, F, Cl, Br, N, OH) in ChEMBL that improve activity, stratified by that of the parent compound. “Potent” denotes parents with activity ≤32 nM, “mid” 32 nM–1 μM, and “weak” 1 μM–1 mM. **b,** For each target in this study, the percentage of analogs that improve or decrease activity by ≥3-fold or ≥10-fold versus their parent. Fold changes were rounded to the nearest integer prior to thresholding (e.g., 9.6–9.9 counted as 10-fold). Error bars denote 95% bootstrap confidence intervals (percentile method; 20,000 resamples). Compounds yielding no measurable curves were counted as ≥10-fold decreases. **c**, Parent-level summary of analog effects in this study (rounded fold-change).

**Extended Data Fig. 2.**
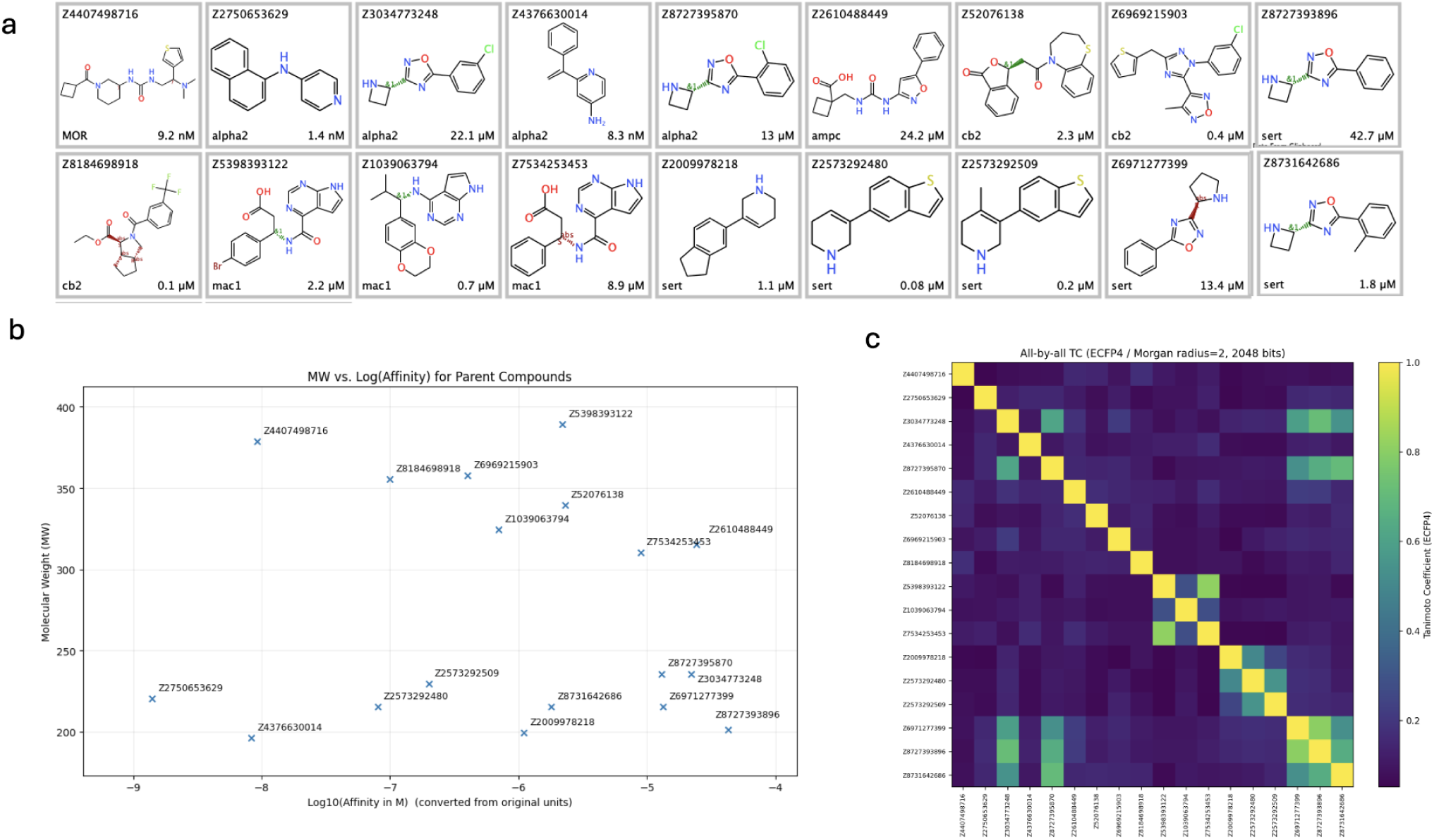
Overview of Parent Compounds’ Structures and Properties. **a.** 2D structures of the parent compounds**. b.** Molecular Weight (MW) and Affinity/potency values for the parent compounds. **c.** Structural Similarity matrix showing the relationships between parent compounds based on molecular fingerprint comparisons.

**Extended Data Fig. 3.**
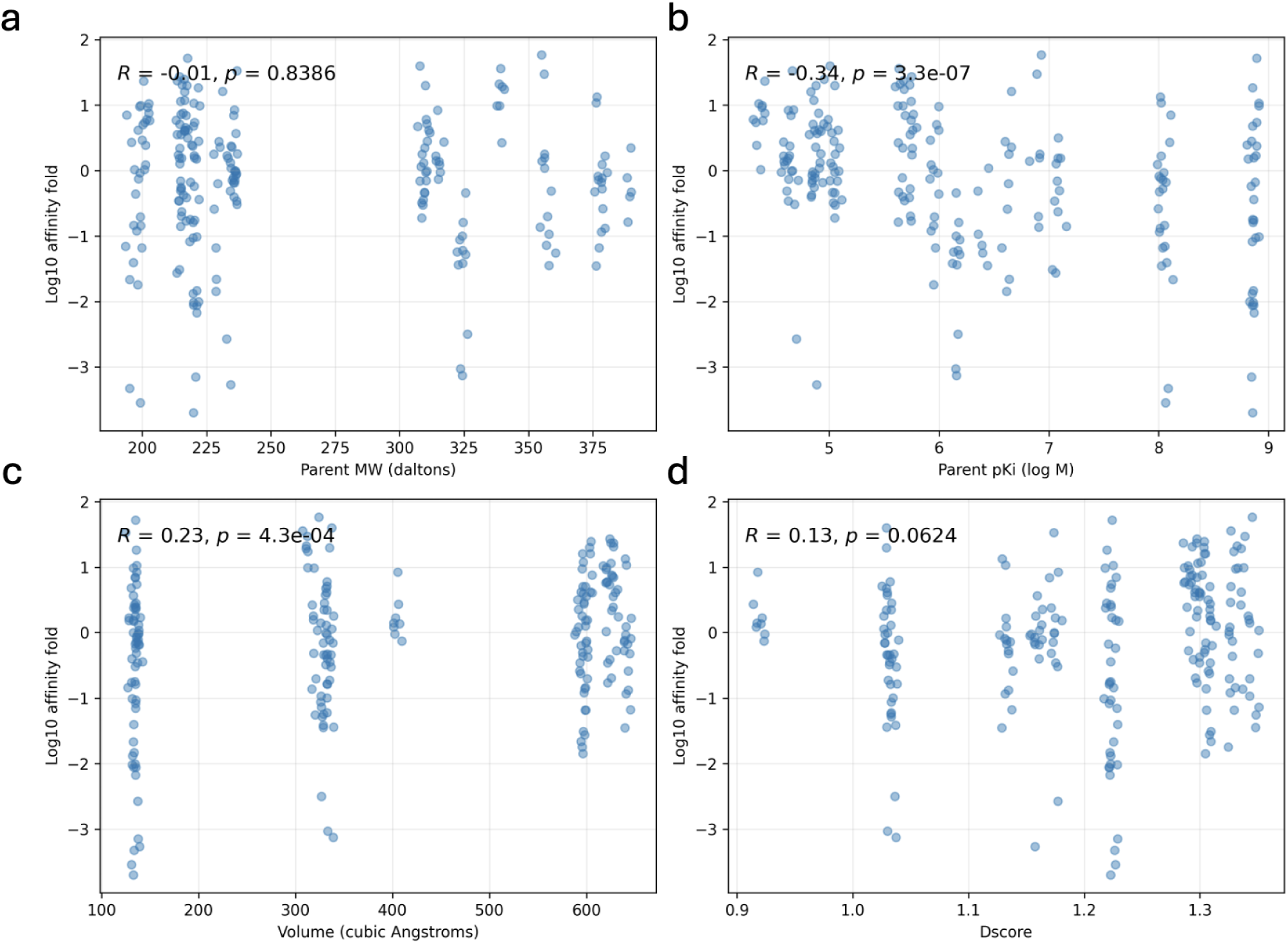
Correlation between parent compound or binding pocket properties and affinity/potency fold change. **a.** Correlation between parent molecular weight (MW) and log₁₀ affinity/potency fold change **b.** Correlation between parent pKi and log₁₀ affinity/potency fold change. **c.** Correlation between binding pocket volume and log₁₀ affinity/potency fold change. **d.** Correlation between SiteMap Dscore and log₁₀ affinity/potency fold change.

**Extended Data Fig. 4.**
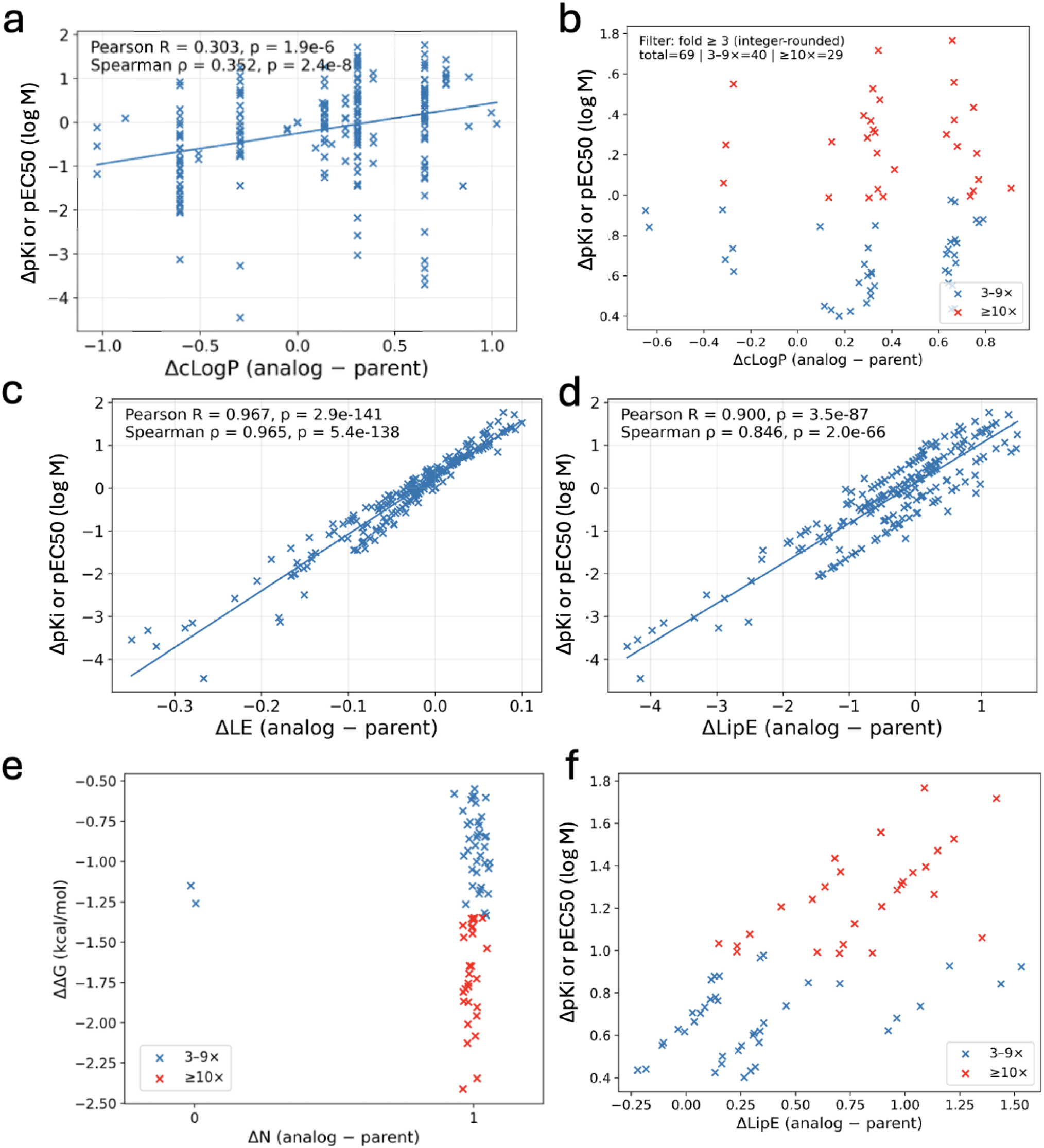
Correlations between physicochemical descriptors and ΔpKi/pEC50 (log M). **a**. Scatter plot of ΔpKi or pEC50 (log M) versus cLogP (RDKit). Each point represents one analog; the solid line indicates a linear fit. Pearson’s R and Spearman’s ρ are shown. **b**. ΔpKi or pEC_50_ (analog − parent) against ΔcLogP for all analog–parent pairs with ≥3× rounded potency/affinity improvement; ≥10× improvements are highlighted in red. **c.** Scatter plot of ΔpKi or pEC50 (log M) versus ΔLE (analog − parent), where LE = 1.37 × pKi or pEC50 / Nheavy. Pearson’s R and Spearman’s ρ are shown. **d.** Scatter plot of ΔpKi or pEC50 (log M) versus ΔLipE (analog − parent), where LipE = pKi or pEC50 − cLogP. Pearson’s R and Spearman’s ρ are shown. **e**. Binding free-energy change versus heavy-atom change for improved analogs. The y-axis shows ΔΔG (kcal/mol), computed from the affinity metric (Ki/IC50/EC50) as ΔG = RT ln(K) (298 K) and ΔΔG = ΔG_analog − ΔG_parent; negative ΔΔG indicates improved binding. The x-axis is ΔN (analog − parent; heavy-atom count difference). Points with rounded fold ≥10 are highlighted in red; rounded 3–9 are shown in blue. **f**. Affinity/potency gains versus changes in lipophilic efficiency. Scatter plot of ΔpKi or pEC50 (analog − parent) against ΔLipE (analog − parent) for all analog–parent pairs with ≥3× rounded potency/affinity improvement; ≥10× improvements are highlighted in red.

**Extended Data Fig. 5.**
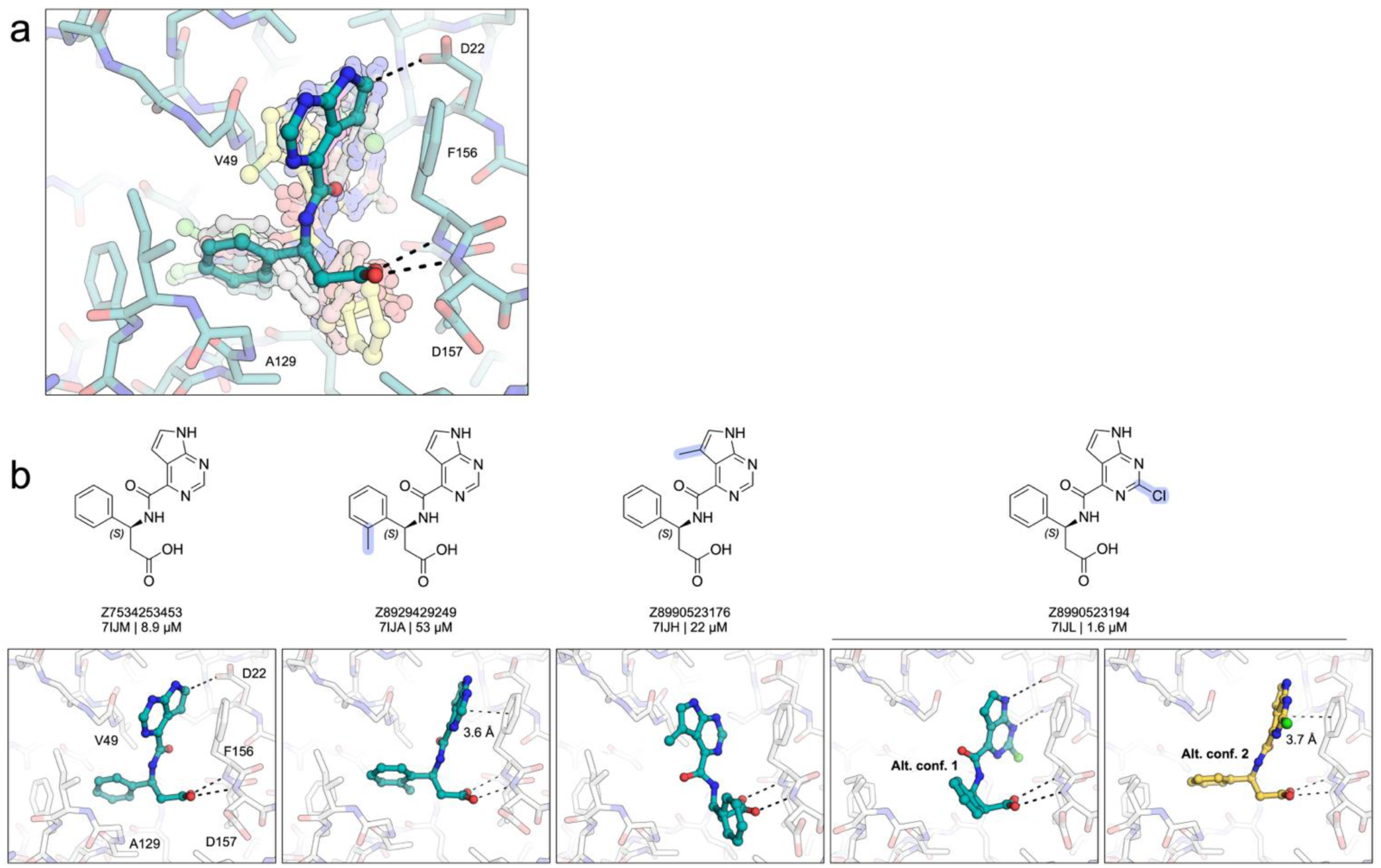
**Structural variability caused by single-atom modifications complicates prediction of affinity**/potency **changes. a.** Overlay of the X-ray crystal structures for 12 ligand-bound complexes from the ‘3453 series showing that single-atom modifications can result in substantial ligand movement or induce multiple binding poses, complicating structure-based analysis and predictions. **b.** Example of a binding pose shift: compounds ‘9249, ‘3176 and ‘3194 adopt markedly different pose compared to the ‘3454 parent. Two conformations were identified for ‘3194.

**Extended Data Fig. 6.**
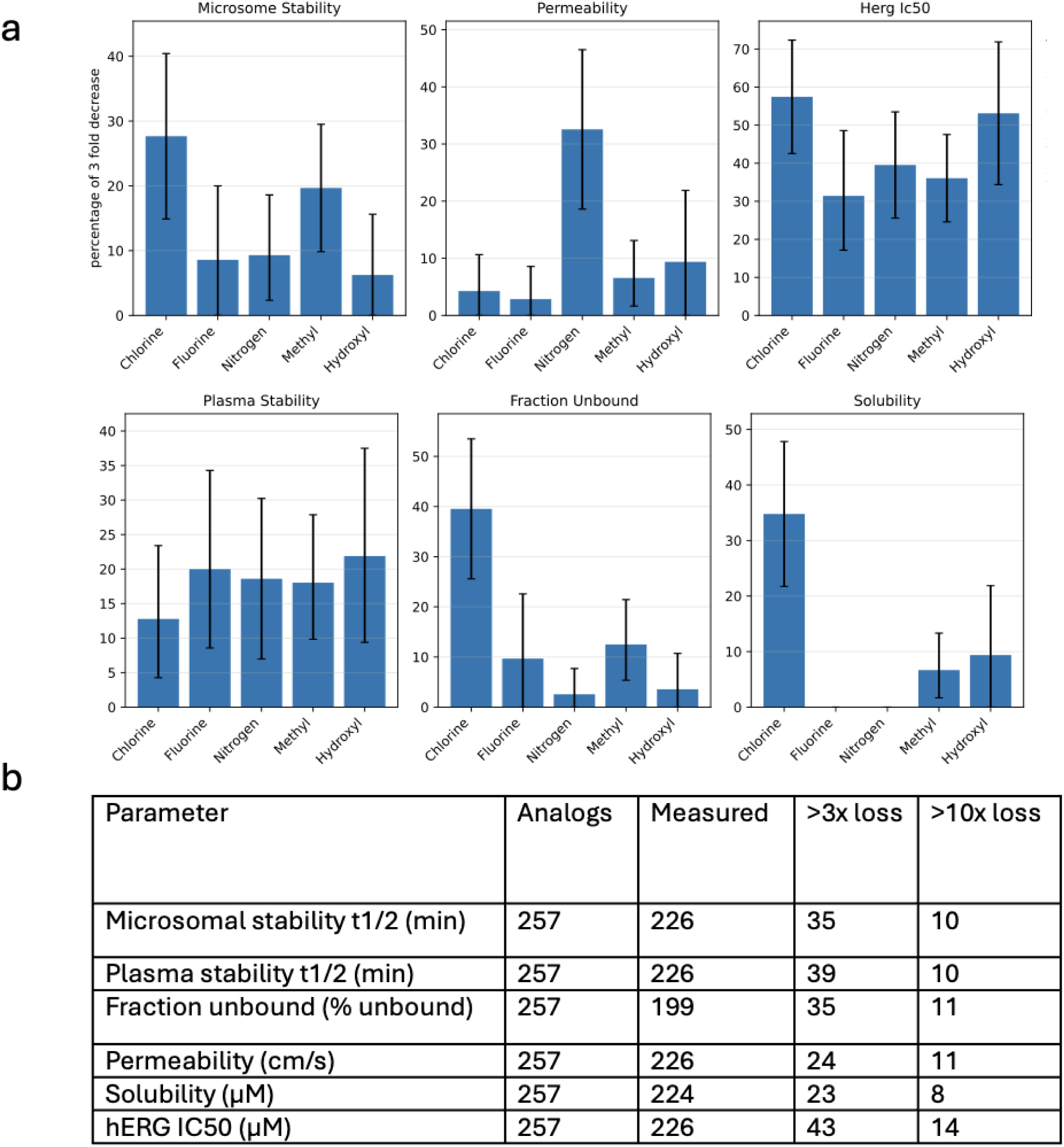
Single-atom substitutions frequently worsen PK properties **a.**Fraction of single-atom substitutions with ≥3-fold decreases in PK properties. Error bars represent 95% confidence intervals, estimated from 10,000 iterations of bootstrap resampling.**b**. Counts of analogs with measured PK parameters and >3-fold or >10-fold losses.

**Extended Data Fig. 7.**
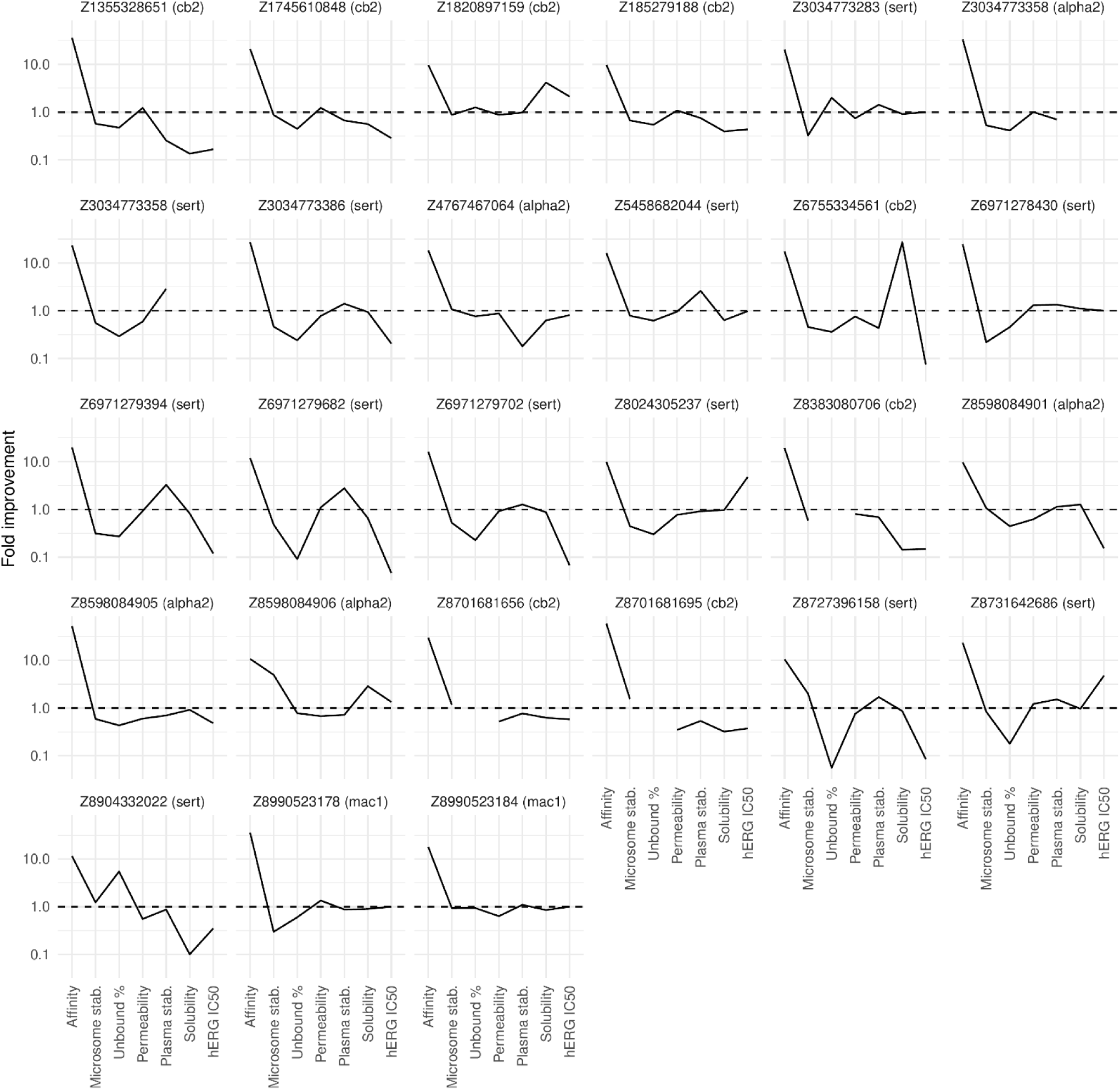
Pharmacokinetic profiles of 29 individual analogs with ≥10-fold affinity/potency improvement. Fold-changes in in vitro pharmacokinetic properties, including microsomal stability, plasma fraction unbound, PAMPA permeability, plasma stability, solubility, and hERG inhibition, for 29 analogs showing ≥10-fold affinity/potency enhancements.

**Extended Data Fig. 8.**
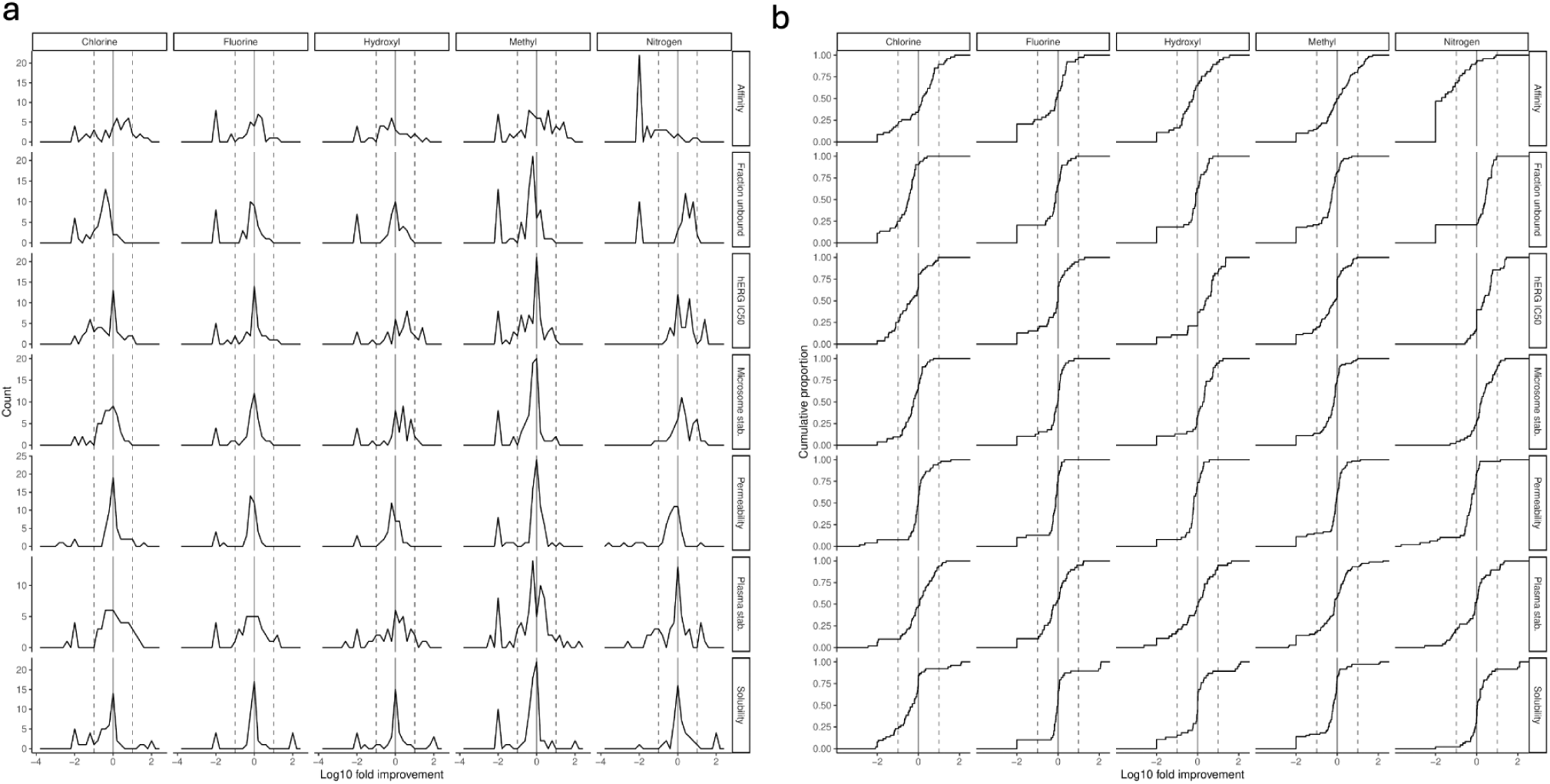
Distribution and cumulative effects of atom modifications on molecular properties. **a.** Density plots showing the distribution of log₁₀ fold improvement for different properties after one-atom modifications (Chlorine, Fluorine, Hydroxyl, Methyl, and Nitrogen). Each row represents a property (e.g., Affinity/potency, Microsome stability, Solubility), and each column corresponds to a specific modification. Vertical dashed lines indicate 0 (no change) and ±0.5 log₁₀ fold change (approximately 3-fold improvement or reduction). **b.** Cumulative distribution plots of the same data as in (a), showing the proportion of analogs achieving various levels of improvement or reduction. This visualization allows for comparison of the shift in property distributions across different modifications.

**Extended Data Table 1.**
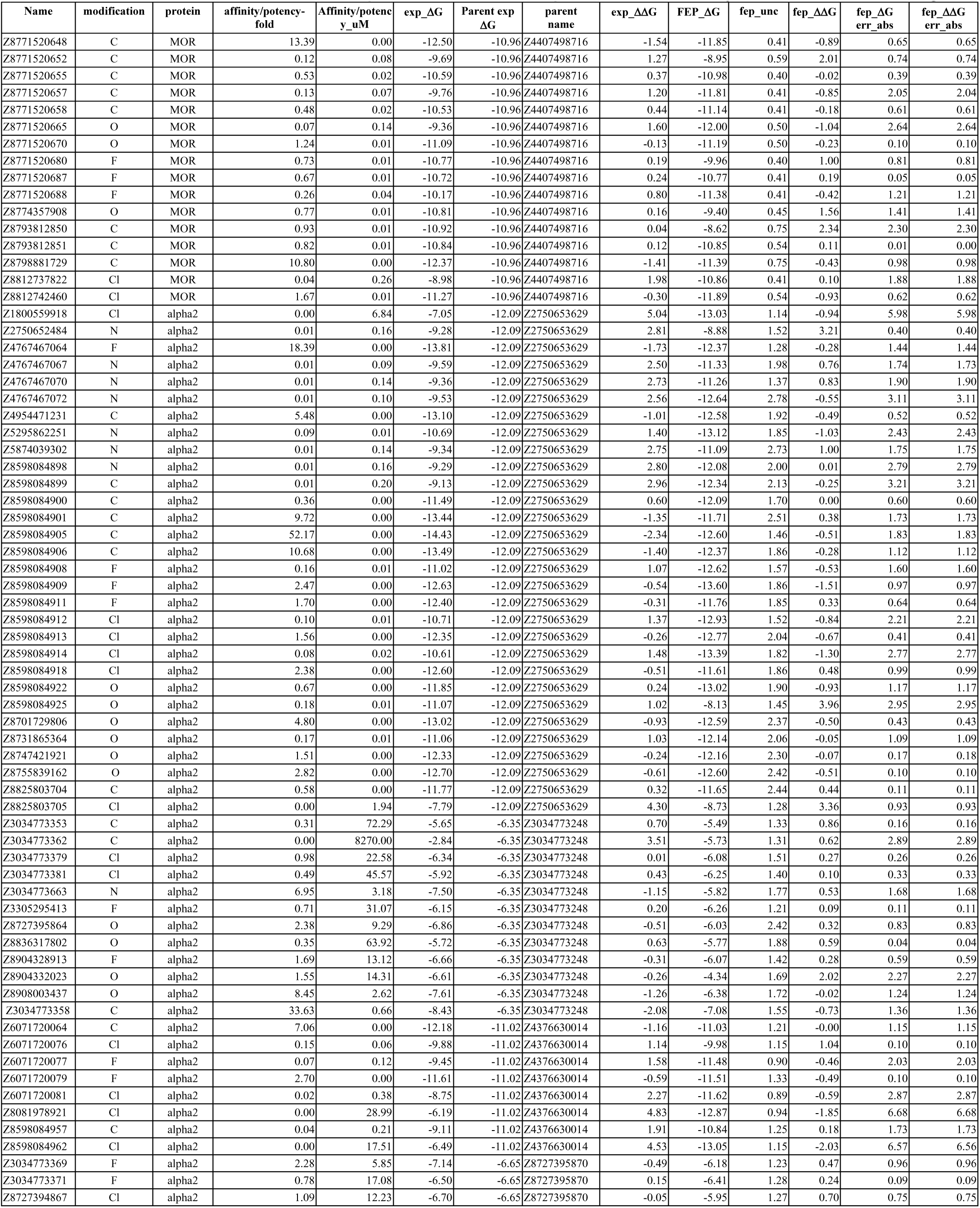
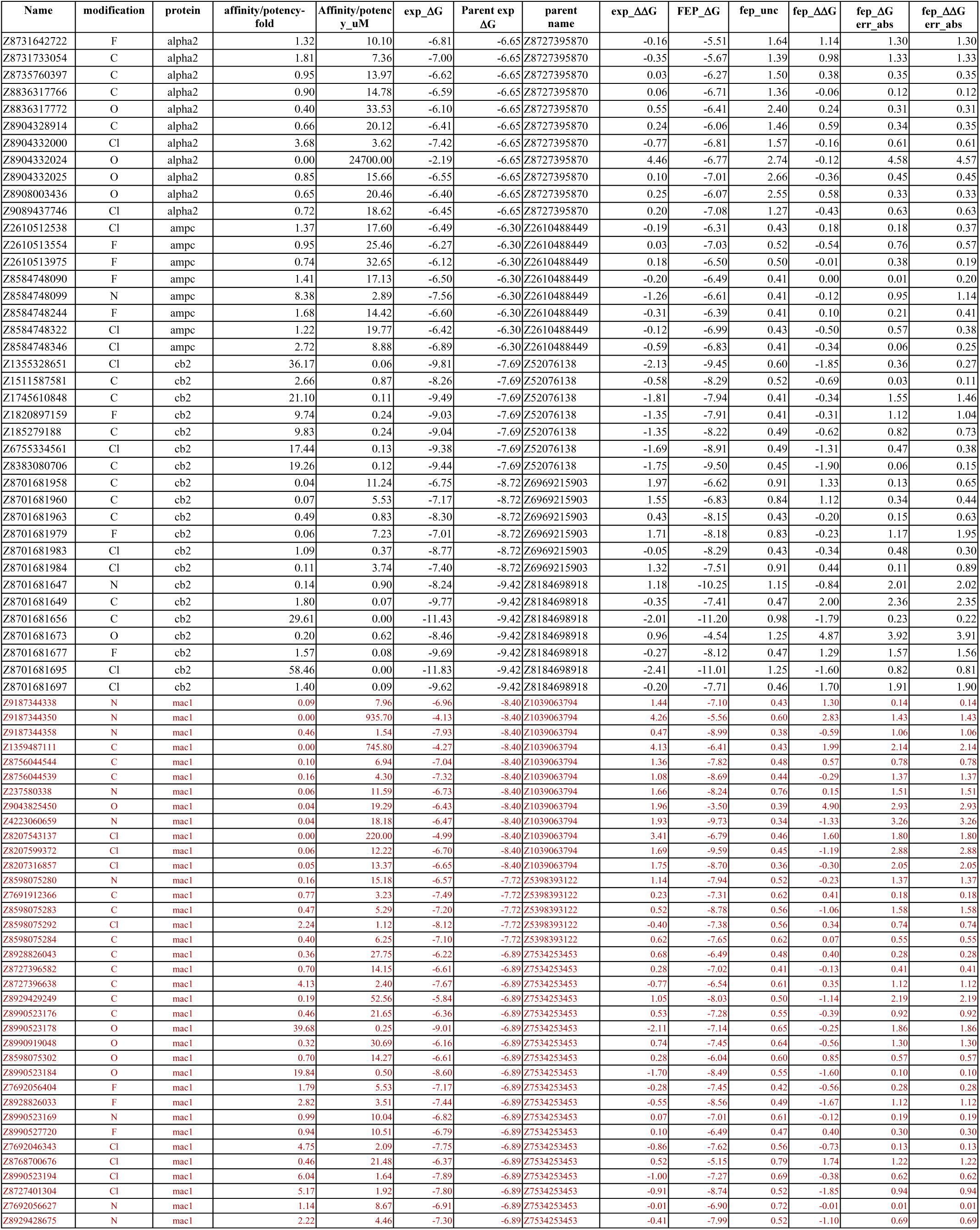
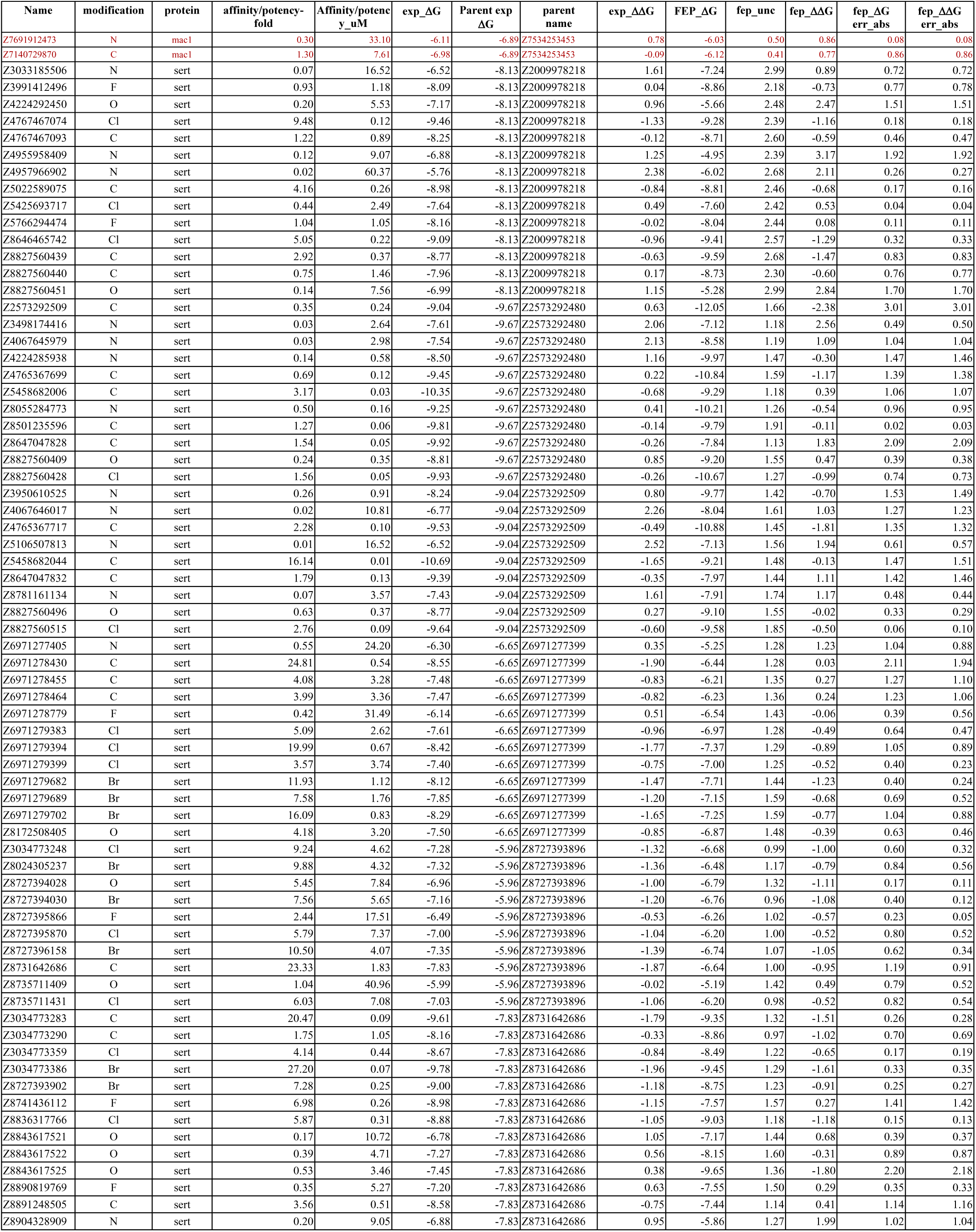
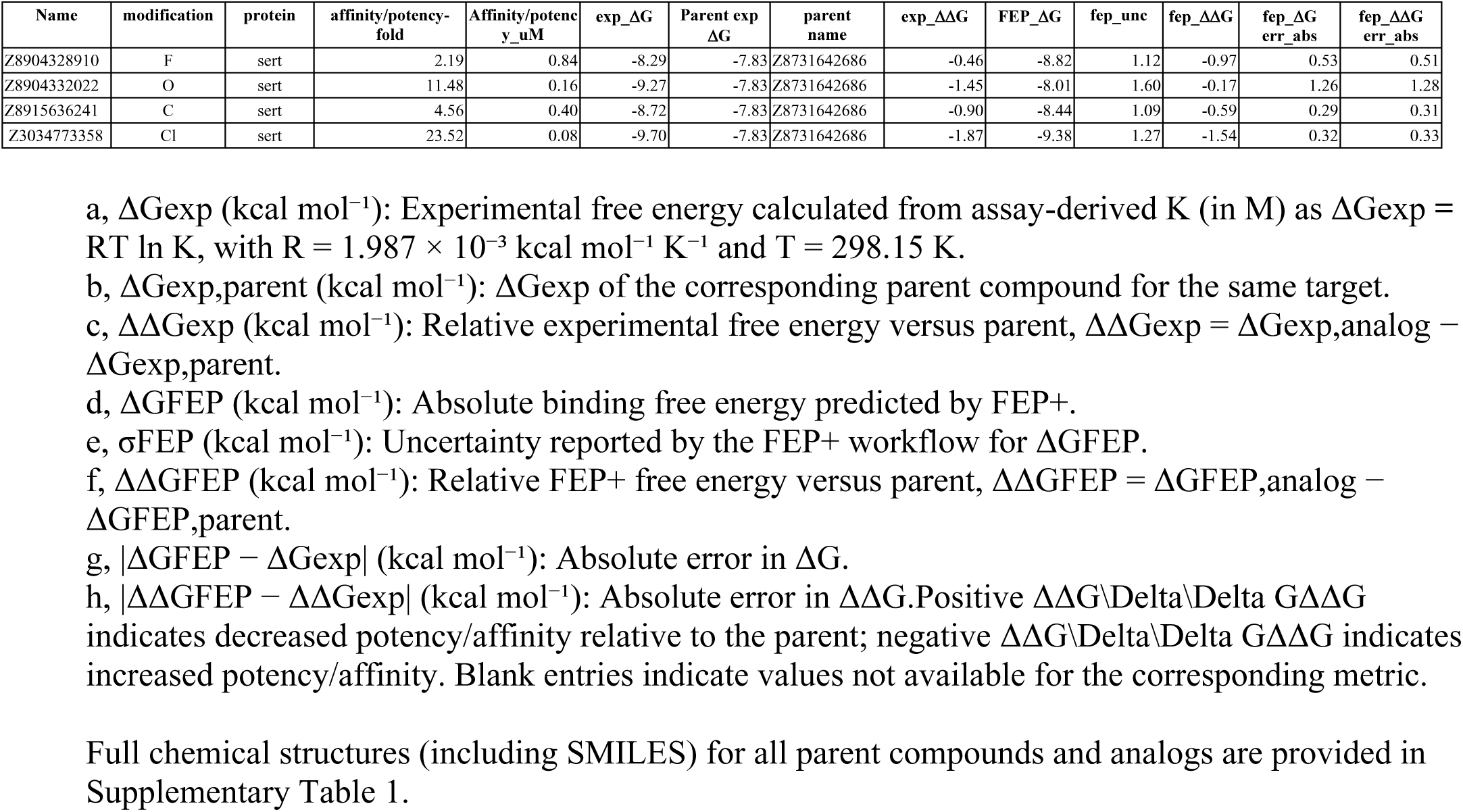
FEP+ predictions and experimental activity readouts for analogs.

**Extended Data Table 2.** The effect of the analogs, relative to parents, on in vitro pharmacokinetics*.

| parent | target | Total compound s | Microsome stability T1/2, min fold change (≥3x) | Plasma stability T1/2, min fold change (≥3x) | ppb % of unbound compound fold change (≥3x) | Solubility, μM fold change (≥3x) | hERG approximate IC50 μM fold change (≥3x) | permeability cm/s fold change (≥3x) |
| --- | --- | --- | --- | --- | --- | --- | --- | --- |
| Z4407498716 | MOR | 16 | 0 | 0 | 0 | 0 | 0 | 0 |
| Z2750653629 | alpha 2 | 31 | 7 | 2 | 4 | 5 | 5 | 0 |
| Z3034773248 | alpha 2 | 18 | 1 | 1 | 3 | 16 | 7 | 0 |
| Z4376630014 | alpha 2 | 10 | 5 | 1 | 7 | 2 | 1 | 0 |
| Z8727395870 | alpha 2 | 18 | 3 | 0 | 6 | 16 | 11 | 0 |
| Z2610488449 | ampc | 10 | 1 | 0 | 2 | 0 | 2 | 4 |
| Z52076138 | cb2 | 7 | 0 | 1 | 0 | 1 | 0 | 0 |
| Z6969215903 | cb2 | 6 | 1 | 2 | 2 | 0 | 0 | 0 |
| Z8184698918 | cb2 | 20 | 1 | 4 | 0 | 0 | 5 | 0 |
| Z1039063794 | mac1 | 12 | 4 | 0 | 1 | 0 | 3 | 0 |
| Z5398393122 | mac1 | 5 | 4 | 1 | 0 | 0 | 0 | 4 |
| Z7534253453 | mac1 | 22 | 1 | 10 | 0 | 0 | 4 | 9 |
| Z2009978218 | sert | 14 | 5 | 4 | 3 | 0 | 2 | 0 |
| Z2573292480 | sert | 11 | 3 | 0 | 3 | 0 | 0 | 0 |
| Z2573292509 | sert | 9 | 3 | 2 | 4 | 6 | 3 | 0 |
| Z6971277399 | sert | 15 | 1 | 3 | 1 | 0 | 1 | 0 |
| Z8727393896 | sert | 14 | 0 | 3 | 0 | 0 | 0 | 0 |
| Z8731642686 | sert | 19 | 0 | 3 | 3 | 0 | 1 | 0 |

\* The MOR parent series and its 16 analogs are not included in this table because they were not profiled for in vitro PK properties.

## References

1 Hughes, J. P., Rees, S., Kalindjian, S. B. & Philpott, K. L. Principles of early drug discovery. Brit J Pharmacol 162, 1239–1249, doi:10.1111/j.1476-5381.2010.01127.x (2011).

2 Bleicher, K. H., Bohm, H. J., Muller, K. & Alanine, A. I. Hit and lead generation: beyond high-throughput screening. Nat Rev Drug Discov 2, 369–378, doi:10.1038/nrd1086 (2003).

3 Paul, S. M. et al. How to improve R&D productivity: the pharmaceutical industry’s grand challenge. Nature Reviews Drug Discovery 9, 203–214, doi:10.1038/nrd3078 (2010).

4 Lipinski, C. A. Lead- and drug-like compounds: the rule-of-five revolution. Drug Discov Today Technol 1, 337–341, doi:10.1016/j.ddtec.2004.11.007 (2004).

5 Stumpfe, D., Hu, H. & Bajorath, J. Evolving Concept of Activity Cliffs. ACS Omega 4, 14360–14368, doi:10.1021/acsomega.9b02221 (2019).

6 Leeson, P. D. & Springthorpe, B. The influence of drug-like concepts on decision-making in medicinal chemistry. Nat Rev Drug Discov 6, 881–890, doi:10.1038/nrd2445 (2007).

7 Wunberg, T. et al. Improving the hit-to-lead process: data-driven assessment of drug-like and lead-like screening hits. Drug Discov Today 11, 175–180, doi:10.1016/S1359-6446(05)03700-1 (2006).

8 Jorgensen, W. L. Efficient drug lead discovery and optimization. Acc Chem Res 42, 724–733, doi:10.1021/ar800236t (2009).

9 Topliss, J. G. Utilization of Operational Schemes for Analog Synthesis in Drug Design. J Med Chem 15, 1006-&, doi:DOI 10.1021/jm00280a002 (1972).

10 Hansch, C. Citation Classic - Rho-Sigma-Pi-Analysis - a Method for the Correlation of Biological-Activity and Chemical-Structure. Cc/Life Sci, 18–18 (1982).

11 Dong, J. et al. ADMETlab: a platform for systematic ADMET evaluation based on a comprehensively collected ADMET database. J Cheminformatics 10, doi:ARTN 29 10.1186/s13321-018-0283-x (2018).

12 Wang, L. et al. Accurate and Reliable Prediction of Relative Ligand Binding Potency in Prospective Drug Discovery by Way of a Modern Free-Energy Calculation Protocol and Force Field. J Am Chem Soc 137, 2695–2703, doi:10.1021/ja512751q (2015).

13 Klimovich, P. V., Shirts, M. R. & Mobley, D. L. Guidelines for the analysis of free energy calculations. J Comput Aided Mol Des 29, 397–411, doi:10.1007/s10822-015-9840-9 (2015).

14 Hajduk, P. J. & Sauer, D. R. Statistical analysis of the effects of common chemical substituents on ligand potency. J Med Chem 51, 553–564, doi:10.1021/jm070838y (2008).

15 Kramer, C., Kalliokoski, T., Gedeck, P. & Vulpetti, A. The Experimental Uncertainty of Heterogeneous Public Data. J Med Chem 55, 5165–5173, doi:10.1021/jm300131x (2012).

16 Booker, T. R., Jackson, B. C. & Keightley, P. D. Detecting positive selection in the genome. Bmc Biol 15, doi:ARTN 98 10.1186/s12915-017-0434-y (2017).

17 Pritchard, J. K. & Cox, N. J. The allelic architecture of human disease genes: common disease - common variant … or not? Hum Mol Genet 11, 2417–2423, doi:DOI 10.1093/hmg/11.20.2417 (2002).

18 Manolio, T. A. et al. Finding the missing heritability of complex diseases. Nature 461, 747–753, doi:10.1038/nature08494 (2009).

19 Fowler, D. M. & Fields, S. Deep mutational scanning: a new style of protein science. Nat Methods 11, 801–807, doi:10.1038/Nmeth.3027 (2014).

20 Wells, J. A. Systematic Mutational Analyses of Protein Protein Interfaces. Method Enzymol 202, 390–411 (1991).

21 Pennington, L. D., Aquila, B. M., Choi, Y., Valiulin, R. A. & Muegge, I. Positional Analogue Scanning: An Effective Strategy for Multiparameter Optimization in Drug Design. J Med Chem 63, 8956–8976, doi:10.1021/acs.jmedchem.9b02092 (2020).

22 Gaulton, A. et al. ChEMBL: a large-scale bioactivity database for drug discovery. Nucleic Acids Res 40, D1100–1107, doi:10.1093/nar/gkr777 (2012).

23 Zdrazil, B. et al. The ChEMBL Database in 2023: a drug discovery platform spanning multiple bioactivity data types and time periods. Nucleic Acids Res 52, D1180–D1192, doi:10.1093/nar/gkad1004 (2024).

24 Muegge, I. & Hu, Y. In Silico Positional Analogue Scanning with Amber GPU-TI. J Chem Inf Model 62, 4448–4459, doi:10.1021/acs.jcim.2c00860 (2022).

25 Schönherr, H. & Cernak, T. Profound Methyl Effects in Drug Discovery and a Call for New C H Methylation Reactions. Angewandte Chemie International Edition 52, 12256–12267 (2013).

26 Cramer, J., Sager, C. P. & Ernst, B. Hydroxyl groups in synthetic and natural-product-derived therapeutics: a perspective on a common functional group. J Med Chem 62, 8915–8930 (2019).

27 Chiodi, D. & Ishihara, Y. “Magic chloro”: profound effects of the chlorine atom in drug discovery. J Med Chem 66, 5305–5331 (2023).

28 Gillis, E. P., Eastman, K. J., Hill, M. D., Donnelly, D. J. & Meanwell, N. A. Applications of fluorine in medicinal chemistry. J Med Chem 58, 8315–8359 (2015).

29 Hopkins, A. L., Groom, C. R. & Alex, A. Ligand efficiency: a useful metric for lead selection. Drug Discov Today 9, 430–431, doi:10.1016/S1359-6446(04)03069-7 (2004).

30 Zhu, T. et al. Hit identification and optimization in virtual screening: practical recommendations based on a critical literature analysis. J Med Chem 56, 6560–6572, doi:10.1021/jm301916b (2013).

31 Kuntz, I. D., Chen, K., Sharp, K. A. & Kollman, P. A. The maximal affinity of ligands. Proc Natl Acad Sci U S A 96, 9997–10002, doi:10.1073/pnas.96.18.9997 (1999).

32 Halgren, T. A. Identifying and Characterizing Binding Sites and Assessing Druggability. J Chem Inf Model 49, 377–389, doi:10.1021/ci800324m (2009).

33 Thomas, M., Bender, A. & de Graaf, C. Integrating structure-based approaches in generative molecular design. Curr Opin Struct Biol 79, 102559, doi:10.1016/j.sbi.2023.102559 (2023).

34 Flowers, J. et al. Expanding automated multiconformer ligand modeling to macrocycles and fragments. Elife 14, doi:10.7554/eLife.103797 (2025).

35 Di, L., Kerns, E. H. & Carter, G. T. Drug-like property concepts in pharmaceutical design. Curr Pharm Des 15, 2184–2194, doi:10.2174/138161209788682479 (2009).

36 Morgan, P. et al. Can the flow of medicines be improved? Fundamental pharmacokinetic and pharmacological principles toward improving Phase II survival. Drug Discov Today 17, 419–424, doi:10.1016/j.drudis.2011.12.020 (2012).

37 Houston, J. B. Utility of in vitro drug metabolism data in predicting in vivo metabolic clearance. Biochem Pharmacol 47, 1469–1479, doi:10.1016/0006-2952(94)90520-7 (1994).

38 Iwatsubo, T. et al. Prediction of in vivo drug metabolism in the human liver from in vitro metabolism data. Pharmacol Ther 73, 147–171, doi:10.1016/s0163-7258(96)00184-2 (1997).

39 Amidon, G. L., Lennernas, H., Shah, V. P. & Crison, J. R. A theoretical basis for a biopharmaceutic drug classification: the correlation of in vitro drug product dissolution and in vivo bioavailability. Pharm Res 12, 413–420, doi:10.1023/a:1016212804288 (1995).

40 Hopkins, A. L., Keseru, G. M., Leeson, P. D., Rees, D. C. & Reynolds, C. H. The role of ligand efficiency metrics in drug discovery. Nat Rev Drug Discov 13, 105–121, doi:10.1038/nrd4163 (2014).

41 Young, R. J. & Leeson, P. D. Mapping the Efficiency and Physicochemical Trajectories of Successful Optimizations. J Med Chem 61, 6421–6467, doi:10.1021/acs.jmedchem.8b00180 (2018).

42 Rocklin, G. J., Mobley, D. L., Dill, K. A. & Hunenberger, P. H. Calculating the binding free energies of charged species based on explicit-solvent simulations employing lattice-sum methods: an accurate correction scheme for electrostatic finite-size effects. J Chem Phys 139, 184103, doi:10.1063/1.4826261 (2013).

43 Kim, H. et al. Artificial Intelligence in Drug Discovery: A Comprehensive Review of Data-driven and Machine Learning Approaches. Biotechnol Bioprocess Eng 25, 895–930, doi:10.1007/s12257-020-0049-y (2020).

44 Kumar, A., Kini, S. G. & Rathi, E. A Recent Appraisal of Artificial Intelligence and In Silico ADMET Prediction in the Early Stages of Drug Discovery. Mini Rev Med Chem 21, 2788–2800, doi:10.2174/1389557521666210401091147 (2021).

45 Fu, L. et al. ADMETlab 3.0: an updated comprehensive online ADMET prediction platform enhanced with broader coverage, improved performance, API functionality and decision support. Nucleic Acids Res 52, W422–W431, doi:10.1093/nar/gkae236 (2024).

46 Myung, Y., de Sa, A. G. C. & Ascher, D. B. Deep-PK: deep learning for small molecule pharmacokinetic and toxicity prediction. Nucleic Acids Res 52, W469–W475, doi:10.1093/nar/gkae254 (2024).

47 Swanson, K. et al. ADMET-AI: a machine learning ADMET platform for evaluation of large-scale chemical libraries. Bioinformatics 40, doi:10.1093/bioinformatics/btae416 (2024).

48 Bergstrom, C. A., Wassvik, C. M., Johansson, K. & Hubatsch, I. Poorly soluble marketed drugs display solvation limited solubility. J Med Chem 50, 5858–5862, doi:10.1021/jm0706416 (2007).

49 Obach, R. S. Prediction of human clearance of twenty-nine drugs from hepatic microsomal intrinsic clearance data: An examination of in vitro half-life approach and nonspecific binding to microsomes. Drug Metab Dispos 27, 1350–1359 (1999).

50. Friden, M., et al. Structure-brain exposure relationships in rat and human using a novel data set of unbound drug concentrations in brain interstitial and cerebrospinal fluids. J Med Chem 52, 6233-6243, doi:10.1021/jm901036q (2009).

51 Fink, E. A. et al. Structure-based discovery of nonopioid analgesics acting through the alpha(2A)-adrenergic receptor. Science 377, eabn7065, doi:10.1126/science.abn7065 (2022).

52 Leung, C. S., Leung, S. S., Tirado-Rives, J. & Jorgensen, W. L. Methyl effects on protein-ligand binding. J Med Chem 55, 4489–4500, doi:10.1021/jm3003697 (2012).

53 Sindt, F., Bret, G. & Rognan, D. On the Difficulty to Rescore Hits from Ultralarge Docking Screens. J Chem Inf Model 65, 5553–5566, doi:10.1021/acs.jcim.5c00730 (2025).

54 Lyu, J. et al. Ultra-large library docking for discovering new chemotypes. Nature 566, 224-+, doi:10.1038/s41586-019-0917-9 (2019).

55 Moroz, Y., Chuprina, A. & Mykytenko, D. Enamine REAL DataBase - an instrumental and practical vehicle for charting new regions of the relevant drug discovery chemical space. Abstr Pap Am Chem S 251 (2016).

56 Verteramo, M. L. et al. Interplay of halogen bonding and solvation in protein-ligand binding. iScience 27, 109636, doi:10.1016/j.isci.2024.109636 (2024).

57 Krimmer, S. G., Betz, M., Heine, A. & Klebe, G. Methyl, ethyl, propyl, butyl: futile but not for water, as the correlation of structure and thermodynamic signature shows in a congeneric series of thermolysin inhibitors. ChemMedChem 9, 833–846, doi:10.1002/cmdc.201400013 (2014).

58. Wermuth, C. G. The Practice of Medicinal Chemistry. (Elsevier Science & Technology, 2008).

59 Gahbauer, S. et al. Iterative computational design and crystallographic screening identifies potent inhibitors targeting the Nsp3 macrodomain of SARS-CoV-2. Proceedings of the National Academy of Sciences 120, e2212931120 (2023).

60 Liu, F. et al. The impact of library size and scale of testing on virtual screening. Nature Chemical Biology, 1–7 (2025).

61 Clopper, C. J. & Pearson, E. S. The use of confidence or fiducial limits illustrated in the case of the binomial. Biometrika 26, 404–413 (1934).

62 Virtanen, P. et al. SciPy 1.0: fundamental algorithms for scientific computing in Python. Nat Methods 17, 261–272 (2020).

63 Lu, C. et al. OPLS4: Improving Force Field Accuracy on Challenging Regimes of Chemical Space. J Chem Theory Comput 17, 4291–4300, doi:10.1021/acs.jctc.1c00302 (2021).

64 Balius, T. E. et al. Testing inhomogeneous solvation theory in structure-based ligand discovery. Proceedings of the National Academy of Sciences 114, E6839–E6846 (2017).

65 Singh, I. et al. Structure-based discovery of conformationally selective inhibitors of the serotonin transporter. Cell 186, 2160–2175. e2117 (2023).

66 Vigneron, S. F. et al. Docking 14 Million Virtual Isoquinuclidines against the μ and κ Opioid Receptors Reveals Dual Antagonists–Inverse Agonists with Reduced Withdrawal Effects. ACS Central Science (2025).

67 Hua, T. et al. Activation and signaling mechanism revealed by cannabinoid receptor-Gi complex structures. Cell 180, 655–665. e618 (2020).

68 Hua, T. et al. Crystal structures of agonist-bound human cannabinoid receptor CB1. Nature 547, 468–471 (2017).

69 Manglik, A. et al. Structure-based discovery of opioid analgesics with reduced side effects. Nature 537, 185–190 (2016).

70 Yung-Chi, C. & Prusoff, W. H. Relationship between the inhibition constant (KI) and the concentration of inhibitor which causes 50 per cent inhibition (I50) of an enzymatic reaction. Biochemical pharmacology 22, 3099–3108 (1973).

71 Schuller, M. et al. Fragment binding to the Nsp3 macrodomain of SARS-CoV-2 identified through crystallographic screening and computational docking. Science advances 7, eabf8711 (2021).

72 Maeda, S. et al. Development of an antibody fragment that stabilizes GPCR/G-protein complexes. Nat Commun 9, 3712, doi:10.1038/s41467-018-06002-w (2018).

73 Goddard, T. D. et al. UCSF ChimeraX: Meeting modern challenges in visualization and analysis. Protein Sci 27, 14–25, doi:10.1002/pro.3235 (2018).

74 Liebschner, D. et al. Macromolecular structure determination using X-rays, neutrons and electrons: recent developments in Phenix. Acta Crystallogr D Struct Biol 75, 861–877, doi:10.1107/S2059798319011471 (2019).

75 Emsley, P., Lohkamp, B., Scott, W. G. & Cowtan, K. Features and development of Coot. Acta Crystallogr D Biol Crystallogr 66, 486–501, doi:10.1107/S0907444910007493 (2010).

76 Schuller, M. et al. Fragment binding to the Nsp3 macrodomain of SARS-CoV-2 identified through crystallographic screening and computational docking. Sci Adv 7, doi:10.1126/sciadv.abf8711 (2021).

77 Correy, G. J. et al. Exploration of structure-activity relationships for the SARS-CoV-2 macrodomain from shape-based fragment linking and active learning. Sci Adv 11, eads7187, doi:10.1126/sciadv.ads7187 (2025).

