## Supplemental figure for "Toward a Random Background for Ligand Optimization"

**Supplementary figure1.Sample size required to constrain the “random background” success rate.**

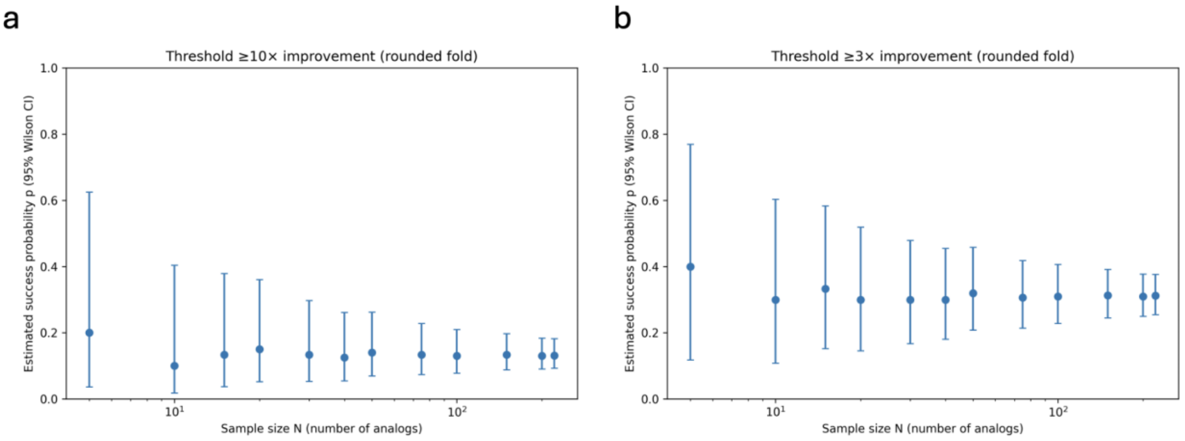


Each analog was treated as a Bernoulli trial for a threshold outcome (success = integer-rounded fold improvement ≥10× in panel a; ≥3× in panel b). Points show the estimated success frequency (p^=k/N) at each sample size N; error bars indicate two-sided 95% binomial confidence intervals computed using the Wilson score method. As N increases, the confidence intervals narrow, showing that pooled analyses with large N provide a well-constrained estimate of the background probability, whereas per-parent sample sizes (typically much smaller) yield wide and often uninformative intervals.

**Supplementary figure 2. dose–response curves for all activity assay**

**SERT parent1 2686**

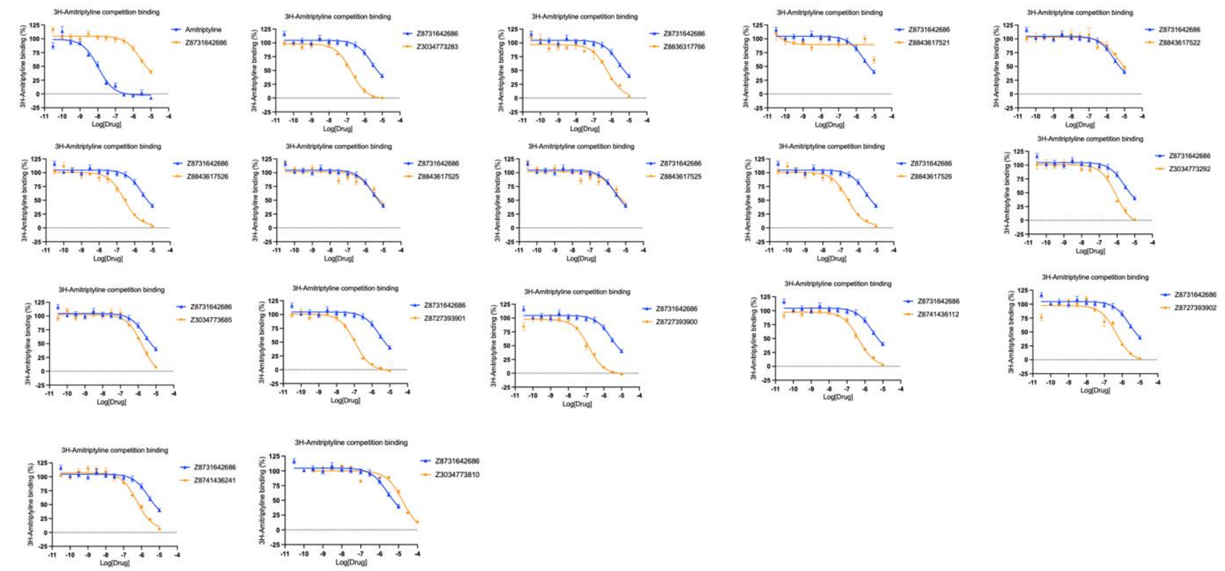

**SERT parent2 2480**

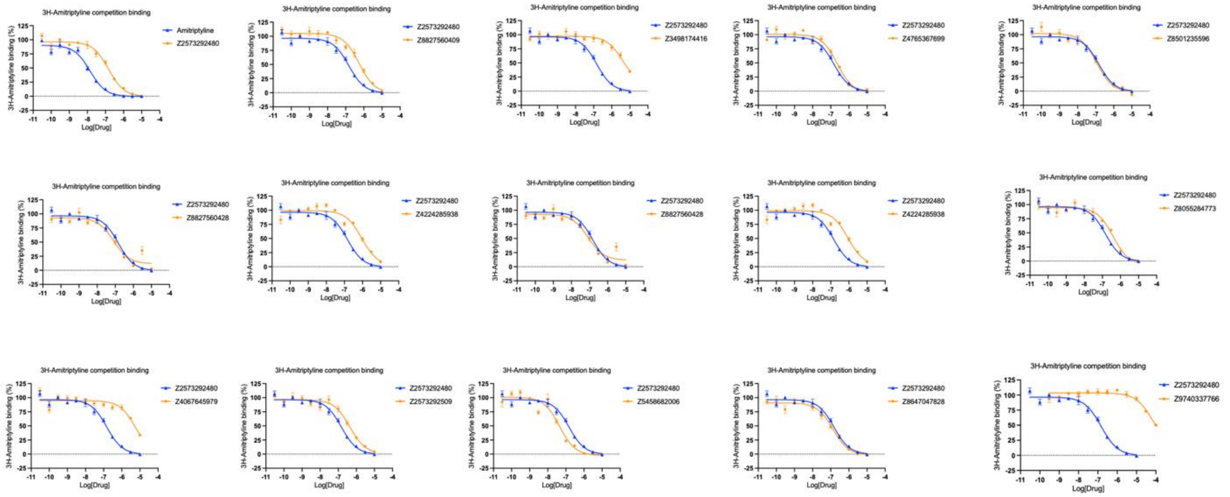

**SERT parent3 8218**

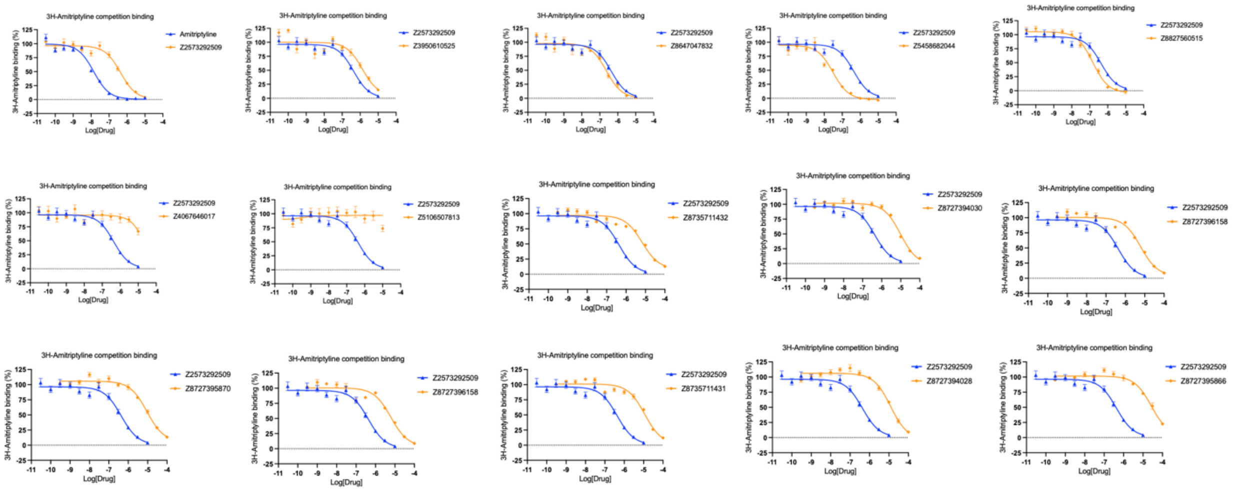


**SERT parent4 2509**

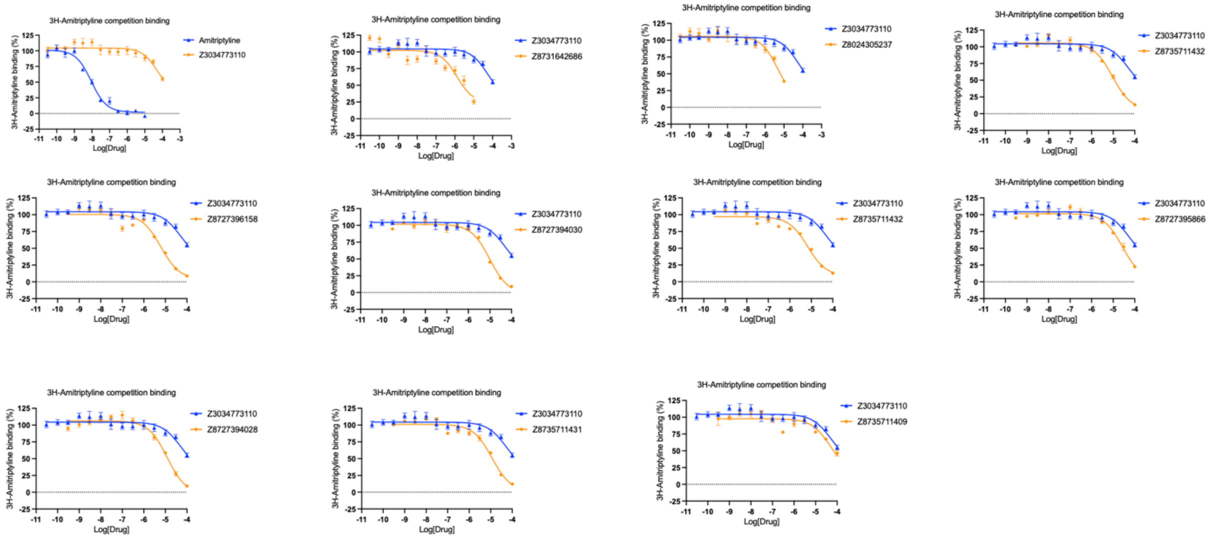

**SERT parent5 3110**

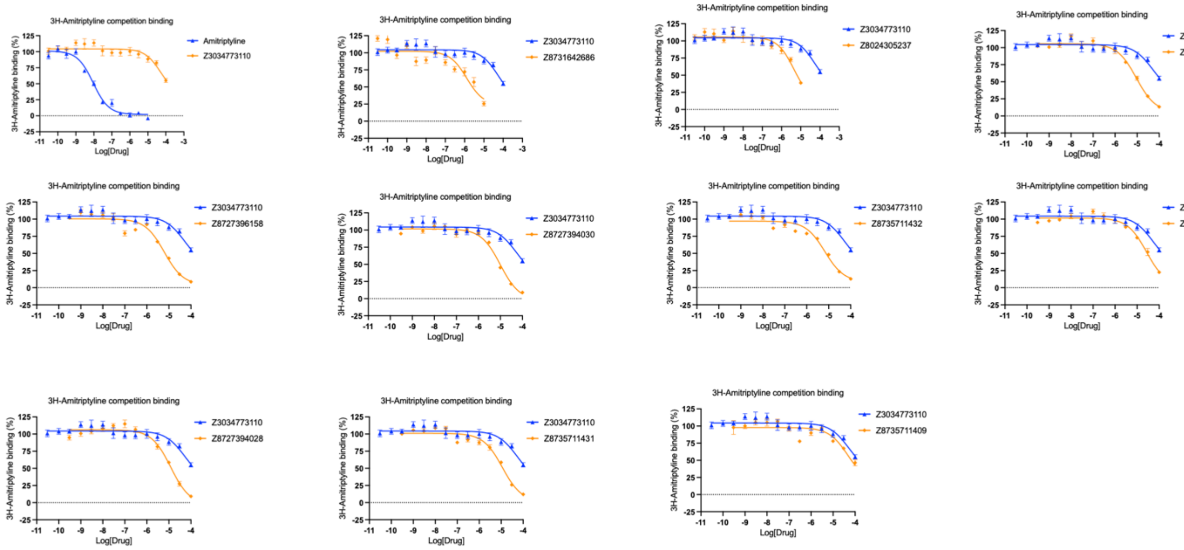


**SERT parent6 7399**

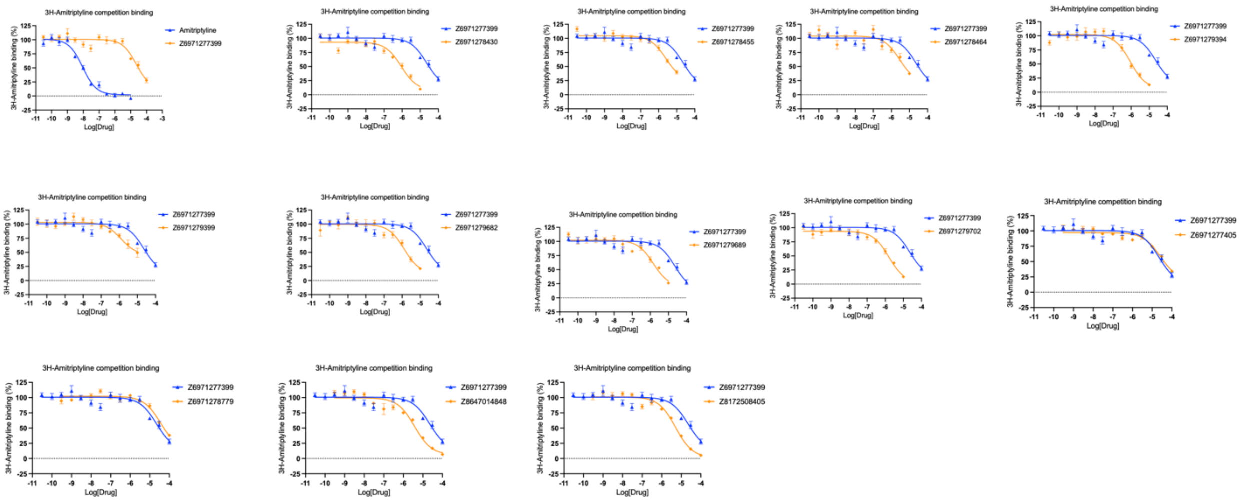

### MOR parent 8716

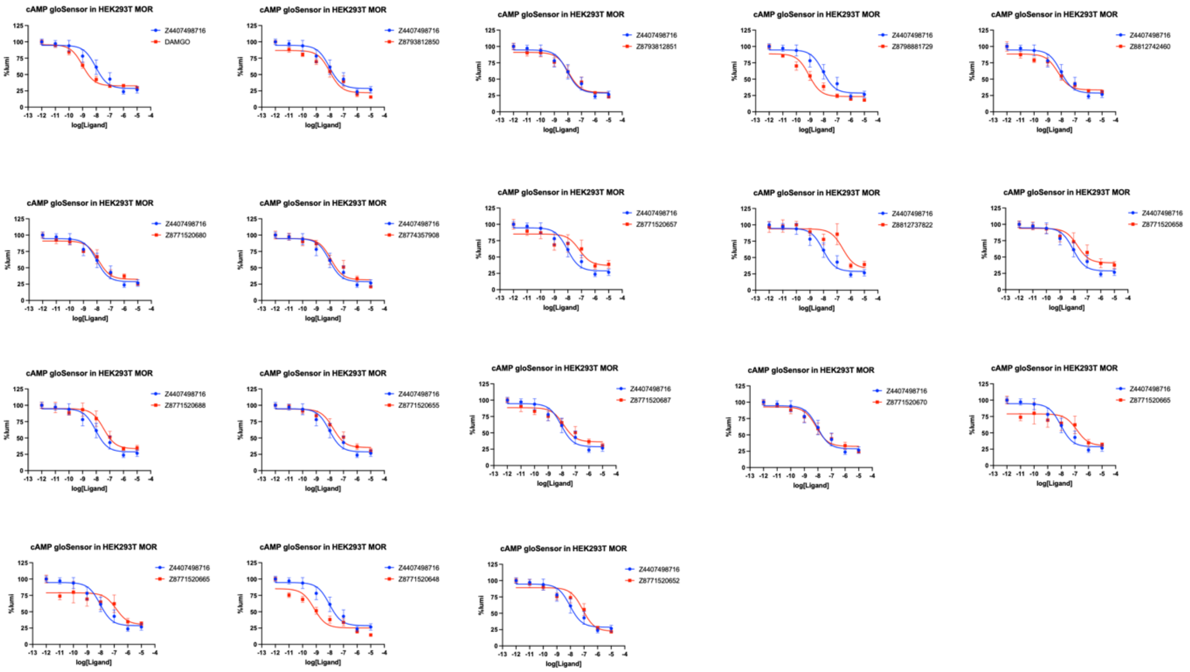


### Alpha2 parent1 3629

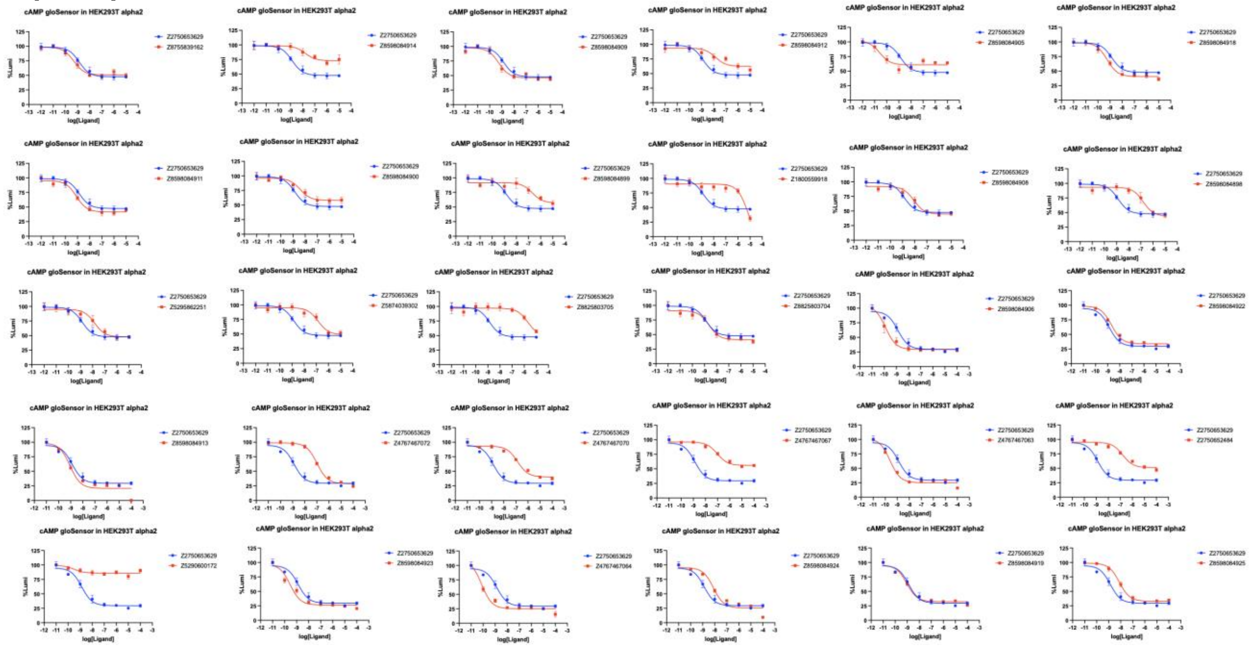

**Alpha2 parent2 0014**

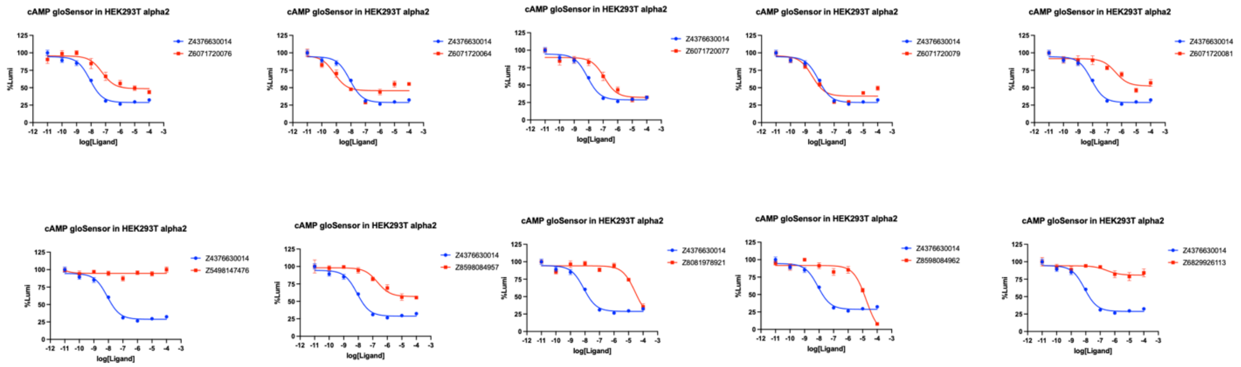


**Alpha2 parent3 3247**

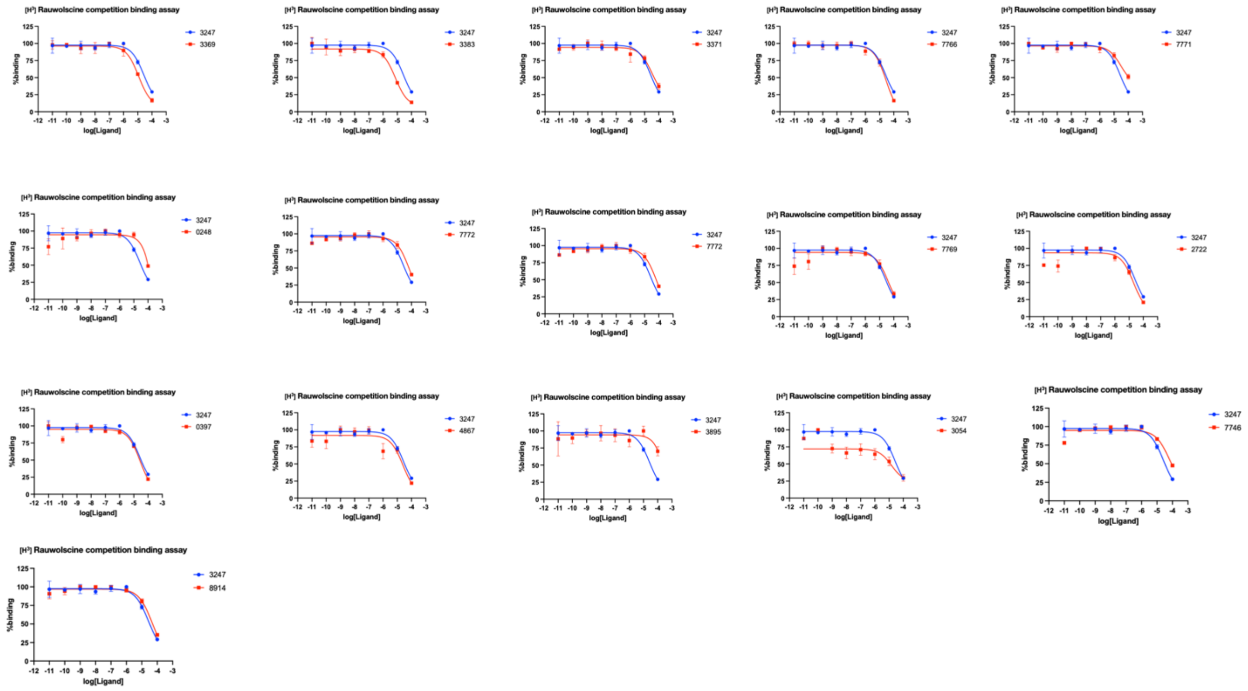

**Alpha2 parent4 3248**

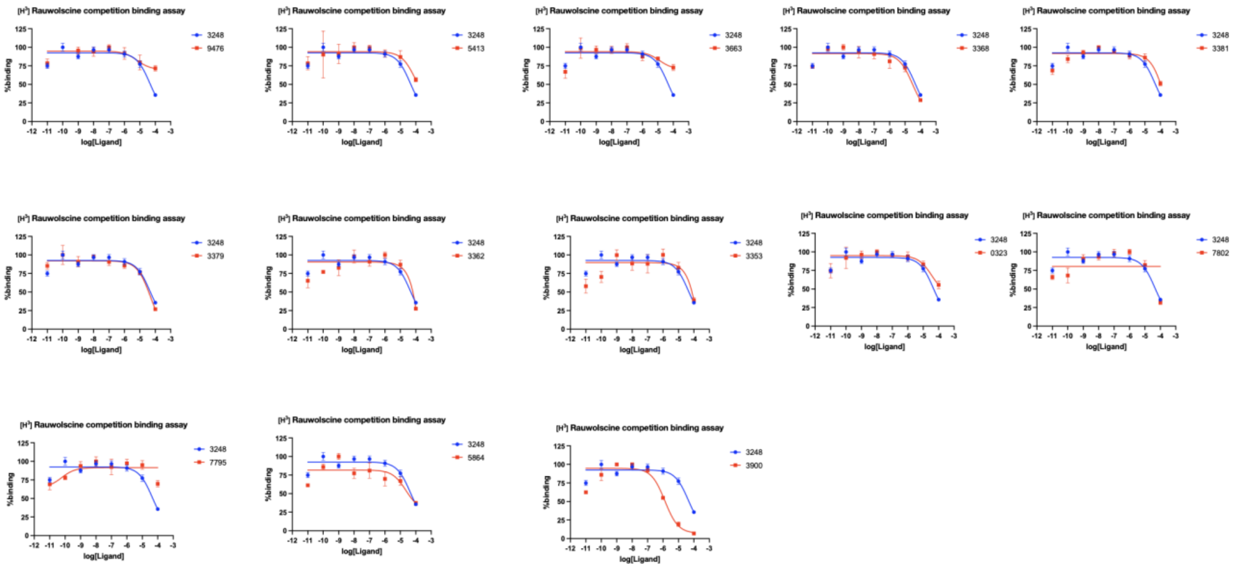


**CB2 parent1 5903**

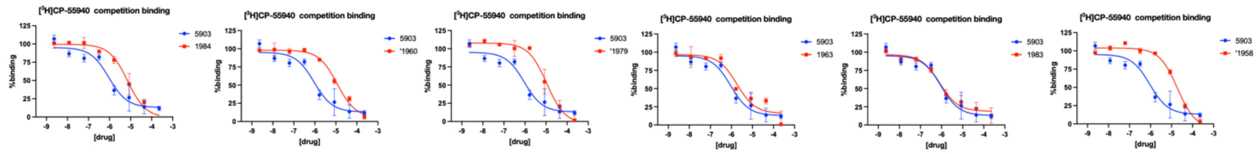


**CB2 parent2 6138**

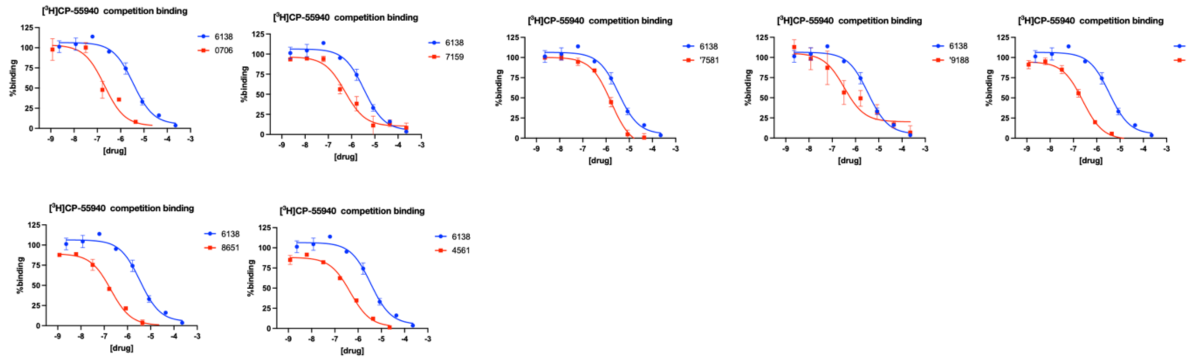

### CB2 parent38918

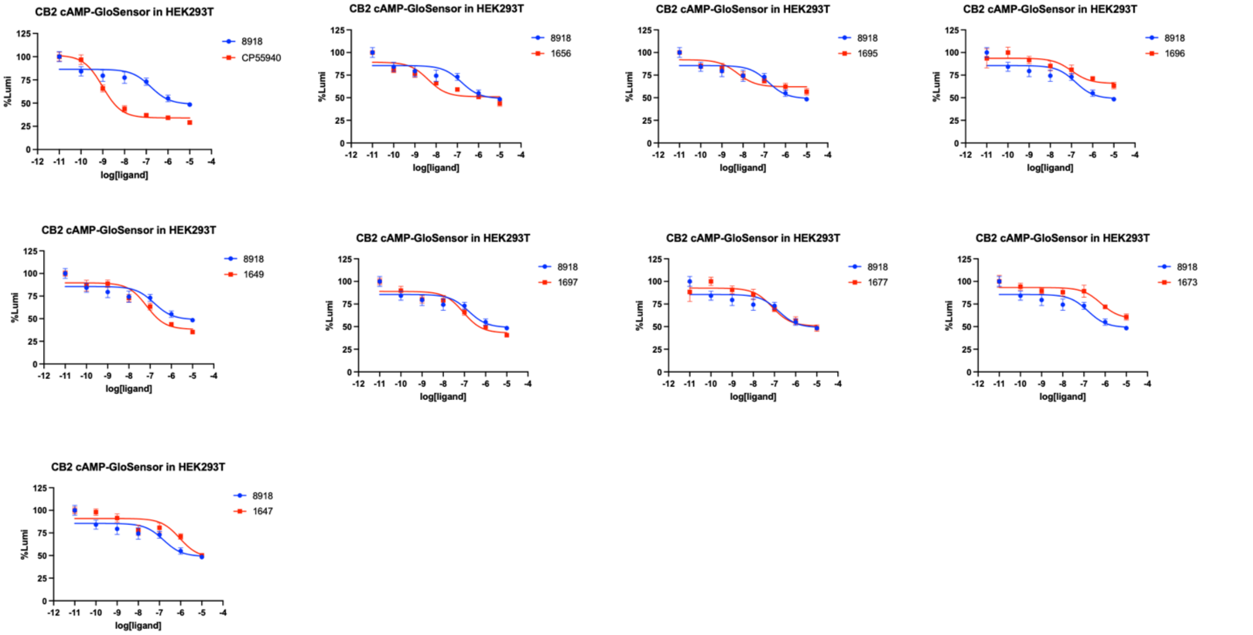


### MAC1 parent1 3122

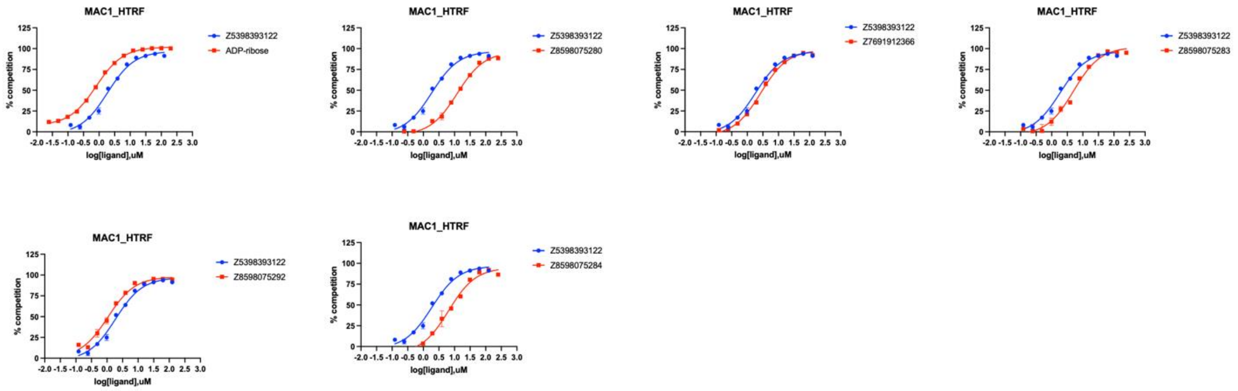

**MAC1 parent2 3453**

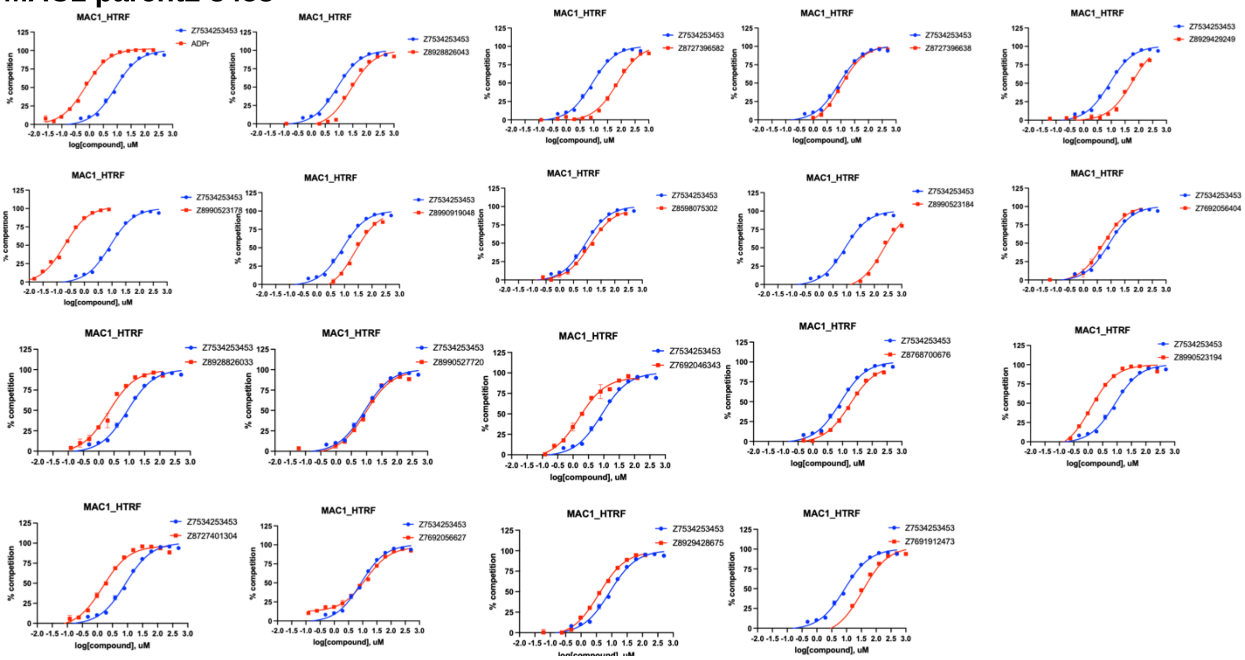


**MAC1 parent3 3794**

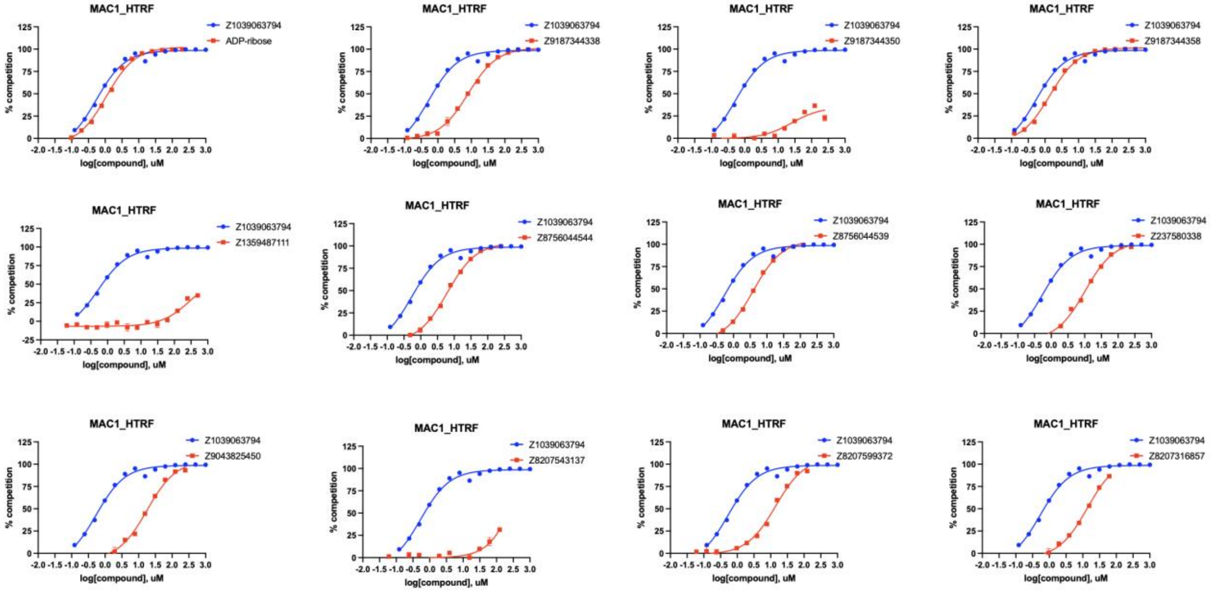

**Supplementary figure 3. Representative concentration–response curves and the corresponding Z values**

a

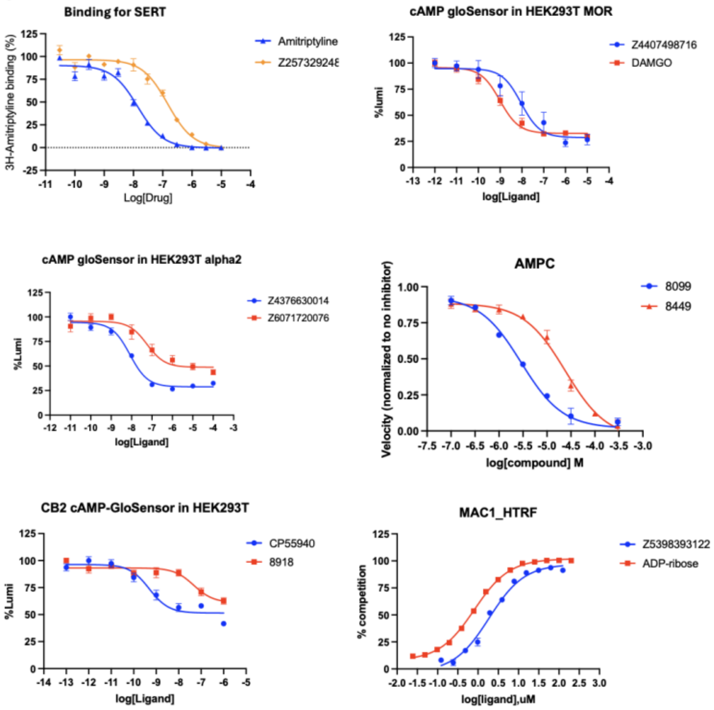


b

| Target | Readout | Parent | Z-value |
|---|---|---|---|
| SERT | Ki | Z2573292480, Z2009978218, Z2573292509, Z6971277399, Z8727393896, Z8731642686 | 0.658 |
| MOR | EC50 | Z4407498716 | 0.5728 |
| alpha2 | EC50 | Z2750653629, Z4376630014 | 0.409 |
| alpha2 | Ki | Z3034773248, Z8727395870 | 0.787 |
| AmpC | Ki | Z2610488449 | 0.493 |
| cb2 | Ki | Z52076138, Z6969215903 | 0.576 |
| cb2 | EC50 | Z8184698918 | 0.571 |
| Mac1 | IC50 | Z5398393122, Z1039063794, Z7534253453 | 0.880 |

a. Representative concentration–response curves for the assays used in this study (SERT; MOR, α2, and CB2; AmpC; Mac1).
b. Corresponding Z-values and the parent compounds used for each target/readout.

**Supplementary figure 4 Receptor expression levels for α2A, CB2, and MOR (relative and absolute measurements)**

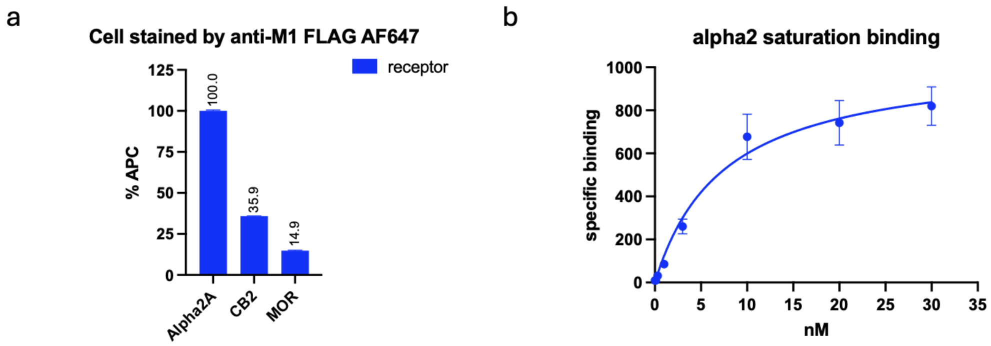


| 96 plate | Plasmid | Expression |
|---|---|---|
| Alpha2A | 100ng | 100 |
| CB2 | 300ng | 35.9 |
| MOR | 600ng | 14.9 |

| | |
|---|---|
| Best-fit values | |
| Bmax | 1046 |
| Kd | 7.466 |
| 95% CI (profile likelihood) | |
| **Bmax** | **870.7 to 1322** |
| Kd | 4.216 to 14.00 |

**a.** Relative quantification of surface receptor expression for α2A, CB2, and MOR by flow cytometry using anti-M1 FLAG–AF647 staining (normalized to α2A = 100%).
**b.** Absolute quantification of α2 receptor expression by radioligand saturation binding, reporting best-fit **Bmax** and **Kd** (with 95% confidence intervals).

**Supplementary figure 5  cAMP responses at different receptor expression levels**

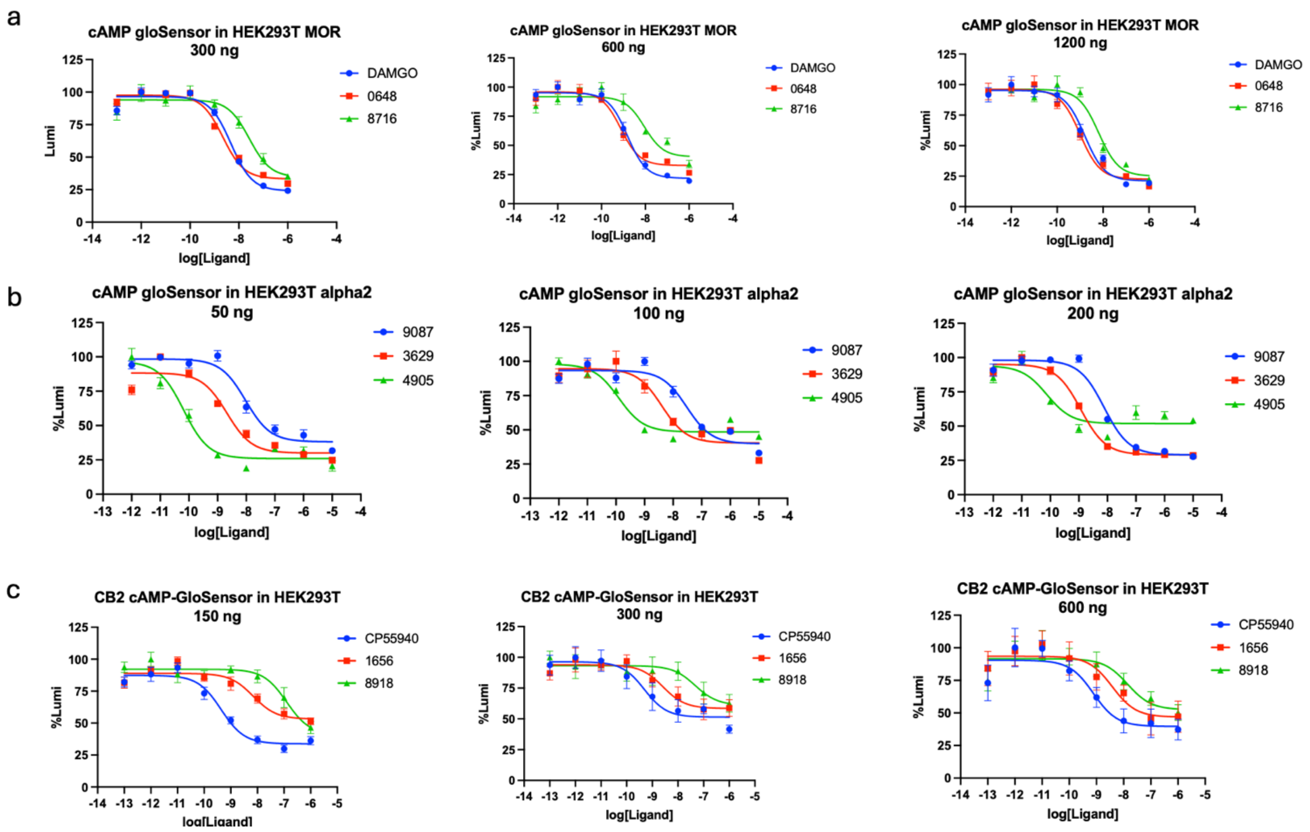


a. MOR cAMP GloSensor concentration–response curves measured in HEK293T cells transfected with increasing amounts of MOR plasmid (300, 600, and 1200 ng), comparing DAMGO and two test ligands.
b. α2AR cAMP GloSensor concentration–response curves measured at increasing receptor plasmid amounts (50, 100, and 200 ng), comparing 9087 and two test ligands.
c. CB2 cAMP GloSensor concentration–response curves measured at increasing receptor plasmid amounts (150, 300, and 600 ng), comparing CP55940 and two test ligands. Points show mean ± error (as plotted) with fitted dose–response curves.

**Supplementary figure 6 PTX-treated cAMP responses and structural rationale for ligand-specific Gs coupling by 4905**

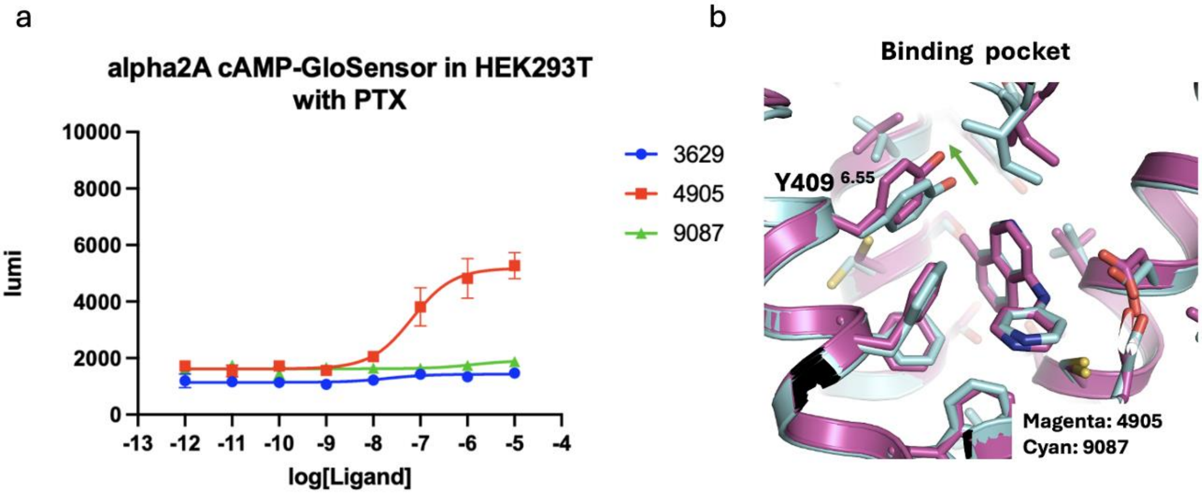


a. Pertussis toxin (PTX)-treated α2A cAMP-GloSensor concentration–response curves in HEK293T cells for 3629, 4905, and 9087. A residual, concentration-dependent increase in cAMP is observed for 4905 at high concentrations, whereas 3629 and 9087 remain flat under PTX treatment.

b. Overlay of the α2A binding pocket comparing 4905 and 9087 complexes (magenta, 4905; cyan, 9087), highlighting an outward shift of Y409^6.55 associated with the additional methyl group in 4905.

**Supplementary figure 7. X-ray crystal structures determined for the SARS-CoV-2 NSP3 Mac1 series ('3122, '3454 and '3794).**

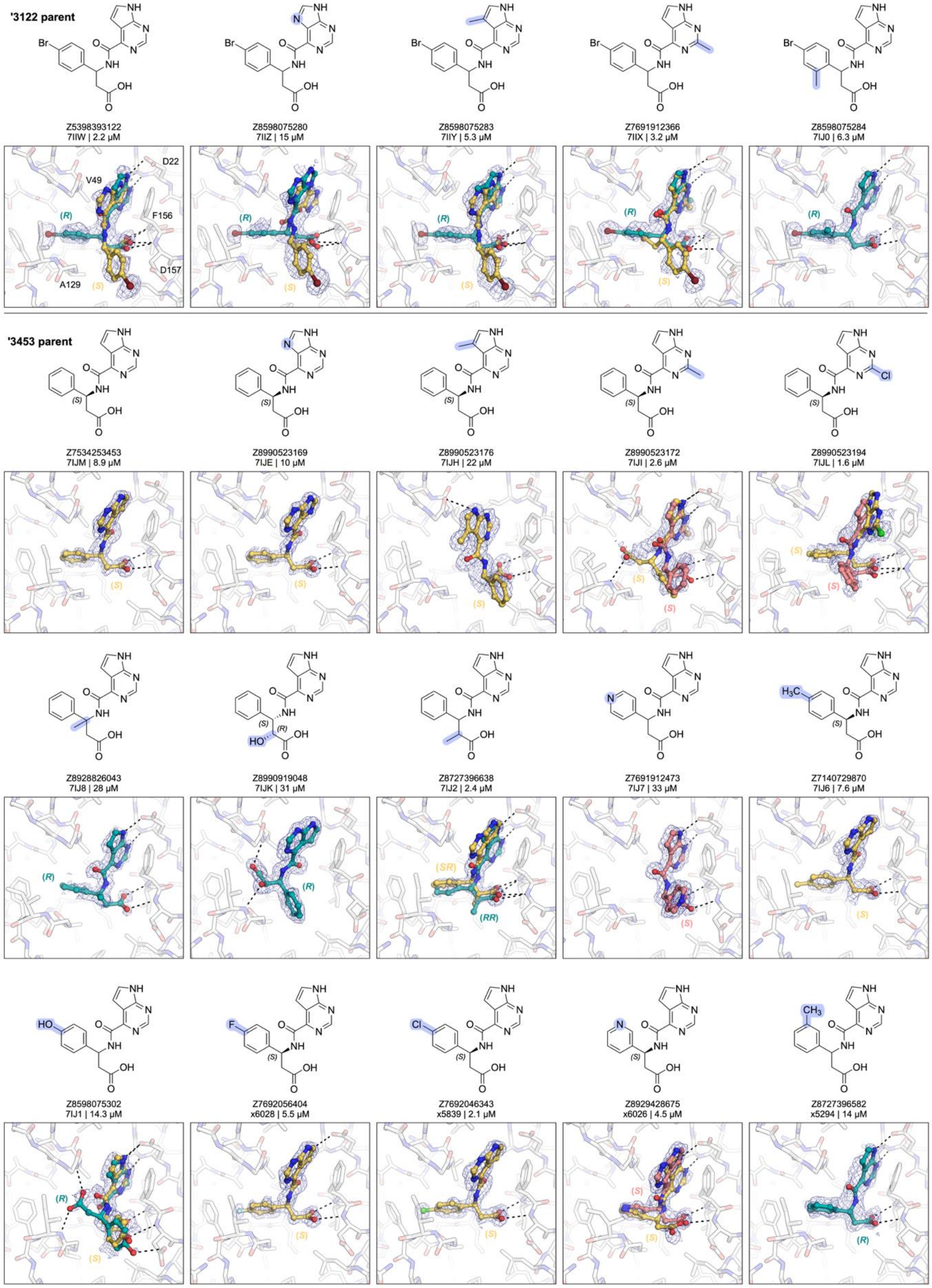

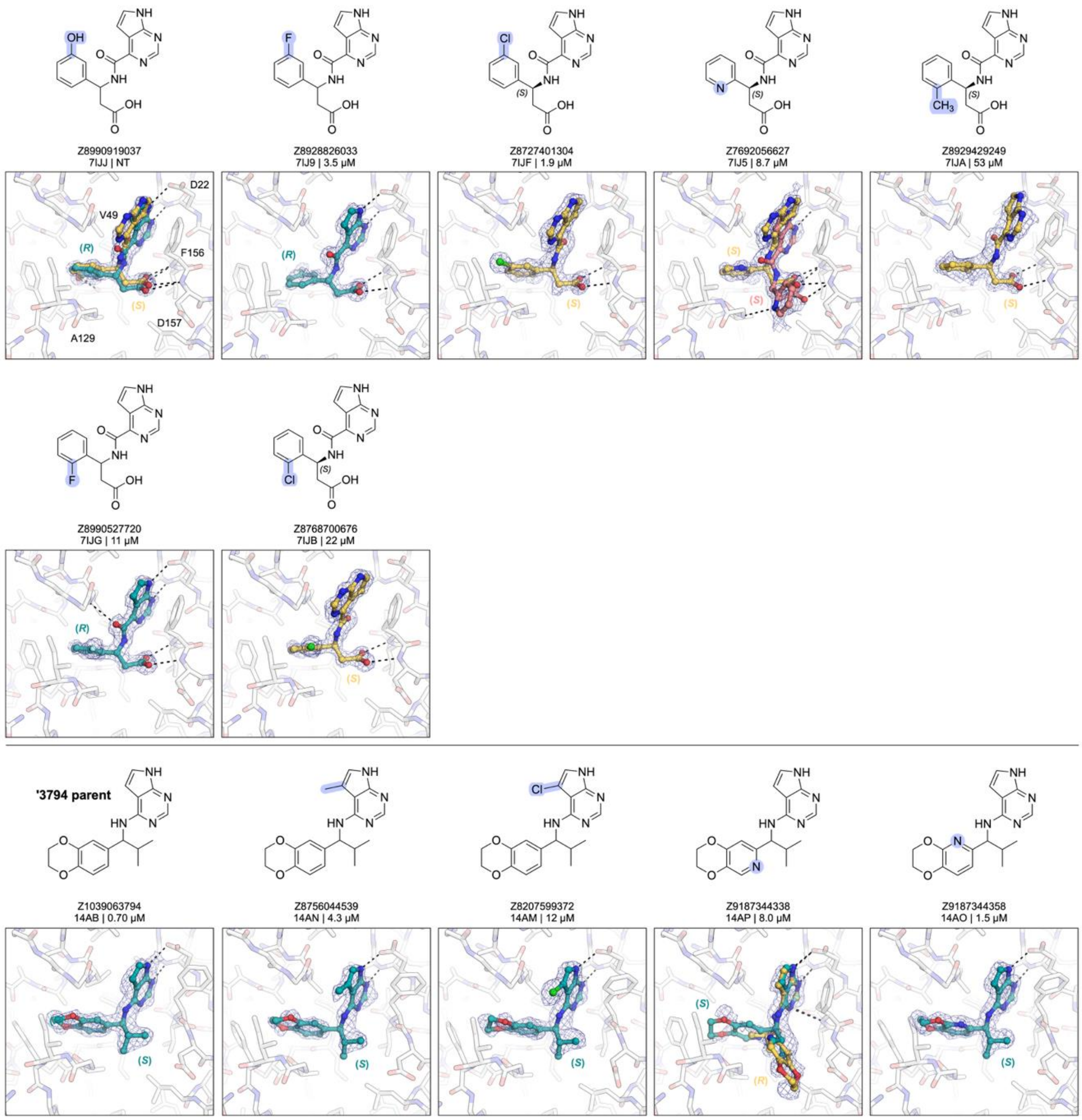


Ligand atoms are colored by isomer and the PanDDA event maps used to identify and model ligands are contoured at 2 σ (blue mesh).
